# Guidelines for extracting biologically relevant context-specific metabolic models using gene expression data

**DOI:** 10.1101/2022.12.04.519052

**Authors:** Saratram Gopalakrishnan, Chintan J. Joshi, Miguel Valderrama Gomez, Elcin Icten, Pablo Rolandi, William Johnson, Cleo Kontoravdi, Nathan E. Lewis

## Abstract

Genome-scale metabolic models comprehensively describe an organism’s metabolism and can be tailored using omics data to model condition-specific physiology. The quality of context-specific models is impacted by (i) choice of algorithm and parameters and (ii) alternate context-specific models that equally explain the -omics data. Here we quantify the influence of alternate optima on microbial and mammalian model extraction using GIMME, iMAT, MBA, and mCADRE. We find that metabolic tasks defining an organism’s phenotype must be explicitly and quantitatively protected. The scope of alternate models is strongly influenced by algorithm choice and the topological properties of the parent genome-scale model with fatty acid metabolism and intracellular metabolite transport contributing much to alternate solutions in all models. mCADRE extracted the most reproducible context-specific models and models generated using MBA had the most alternate solutions. There were fewer qualitatively different solutions generated by GIMME in *E. coli*, but these increased substantially in the mammalian models. Screening ensembles using a receiver operating characteristic plot identified the best-performing models. A comprehensive evaluation of models extracted using combinations of extraction methods and expression thresholds revealed that GIMME generated the best-performing models in *E. coli*, whereas mCADRE is better suited for complex mammalian models. These findings suggest guidelines for benchmarking -omics integration algorithms and motivate the development of a systematic workflow to enumerate alternate models and extract biologically relevant context-specific models.

## 1. INTRODUCTION

The physiological state of a cell is mediated by an intricate network of signaling pathways, gene regulatory networks and metabolic reactions. Gene expression data provide functional insights into the modulation of cellular phenotype (Manzoni et al., 2018), biological features of disease states (Borrageiro et al., 2018; Dickson, 2021; Kori and Yalcin Arga, 2018; Pedrotty et al., 2012), cellular differentiation and tissue-specific functions (Burke et al., 2020; Uhlen et al., 2016; Watcham et al., 2019), and cellular responses to environmental perturbations (Kochanowski et al., 2017). Although many tools improve the coverage of gene expression data analysis, to gain more functional insights into the modulation of cell state (Nguyen et al., 2019), quantitative assessments using genome-scale models (GEMs) can provide rich mechanistic insights.

GEMs are a comprehensive repository of biochemical reactions encoded within the genome of an organism (Gu et al., 2019) that reflect its metabolic capabilities. The sheer size (e.g., number of reactions) of eukaryotic genome-scale models introduces computational and data availability bottlenecks to parameterize quantitative integration techniques such as whole-cell modeling (Macklin et al., 2020), ME-Models (O’Brien et al., 2013), or kinetic models (Gopalakrishnan et al., 2020; Khodayari and Maranas, 2016). The integration of transcriptomics with GEMs has been invaluable to the scientific community for nearly two decades (Blazier and Papin, 2012; Robaina Estevez and Nikoloski, 2014). For example, transcriptomics data can be integrated with eukaryotic models through binarization of enzyme abundance levels to “ON” or “OFF” states after thresholding associated gene expression levels and evaluating gene-protein-reaction (GPR) relationships to yield context-specific models that represent the condition-specific metabolism of the organism. However, inactivating reactions based on thresholding alone leads to fragmented metabolic networks that are incapable of predicting any meaningful flux distributions (hereafter known as flux inconsistent networks) (Åkesson et al., 2004). Flux consistency must be restored using gap-filling algorithms, which seek to preserve the validity of the model. Several algorithms have been developed over the past decade, each with its own unique approach for extracting flux-consistent sub-networks. Context-specific models generated using various model extraction methods have been previously applied to study human tissue-specific metabolism (Jerby et al., 2010), identify biomarkers in NAFLD (Mardinoglu et al., 2014), cancer (Zielinski et al., 2017), and diabetes (Bordbar et al., 2011; Kumar et al., 2014), propose potential anti-cancer drug targets (Pacheco et al., 2019), and optimize bioprocessing for drug manufacturing (Fouladiha et al., 2020; Schinn et al., 2021a).

Model extraction methods are broadly classified into optimization-based and pruning-based methods. Optimization-based methods are broadly classified into the GIMME-like family of methods (Becker and Palsson, 2008) and the iMAT-like family of methods (including iMAT (Zur et al., 2010), INIT (Agren et al., 2012), and tINIT (Agren et al., 2014)) and rely on solving a linear or mixed-integer programming problem to extract context-specific models. The objective varies based on the method and generally maximizes removal of poorly expressed genes (as in the GIMME-like methods) or inclusion of highly expressed genes (as in iMAT and INIT) and may enforce minimum flux through certain required phenotype-defining pathways (also known as required metabolic functions (RMFs)) as implemented in tINIT. On the other hand, pruning-based methods like MBA (Jerby et al., 2010), FASTCORE (Vlassis et al., 2014), mCADRE (Wang et al., 2012), and CORDA (Schultz and Qutub, 2016) extract context-specific models by first identifying a candidate list of reactions to be removed and then pruning the genome-scale models one reaction at a time, until no more reactions can be removed without losing information about the cell’s phenotype. While optimization-based methods are faster and better at protecting flux through known metabolic functions, pruning-based methods allow evidence-based retention of reactions, thereby generating models that are more representative of the physiological state being investigated (Robaina Estevez and Nikoloski, 2014).

The content and quality of an extracted model depends on the choice of model extraction method, the threshold applied to gene expression data to identify active and inactive reactions, and the coverage of data. Previous studies (Opdam et al., 2017; Richelle et al., 2019b) revealed the choice of method and the threshold strongly influencing model content. However, an overlooked factor influencing model content is whether model extraction methods yield a unique context-specific model. Alternate optimal solutions arise when there are multiple combinations of reactions associated with poorly expressed genes that can be retained to restore flux consistency of the metabolic network but cannot be effectively resolved using the available gene expression data. Typically, these include isozymes utilizing different cofactors (e.g., NAD vs NADP) and alternate biosynthetic routes. The scope and disparity of alternate optimal solutions is a measure of reproducibility of each model extraction algorithm and sufficiency of data. To account for alternate optimal solutions, the algorithm EXAMO first identifies all fluxes that are active in all alternate solutions generated by iMAT and uses this set of reactions as high-confidence reactions in MBA (Rossell et al., 2013). Robaina-Estevez and Nikoloski (2017) developed a framework to quantify alternate optima in flux-centric extraction methods such as RegrEx and CORDA and revealed that the variability in extracted model topology stemmed from different combinations of 58% of the reactions that were flagged for removal. Therefore, it is necessary to identify and quantify the variability in extracted context-specific models and screen potential alternate solutions using appropriate data (gene knockout data, fluxomics, endo-metabolomics, etc.) so that extracted models are sufficiently accurate to identify meaningful intervention strategies for therapeutic design or metabolic engineering applications of interest. In addition, a framework to enumerate and screen the space of alternate solutions will provide insights into the reproducibility of existing model extraction algorithms and establish a platform to benchmark future omics-integration algorithms.

This study comprehensively assesses the importance of quantitatively protecting flux through RMF reactions (the biomass production reaction, in this case) and the effect of choice of threshold and extraction method on the scope of alternate optimal solutions during transcriptomics-based model extraction in *E. coli*, CHO-S, and a renal cancer cell line (786O). Ensembles of 100 context-specific models were extracted using combinations of parameters selected from five thresholding approaches (global 80^th^ percentile, global 75^th^ percentile, global 60^th^ percentile, StanDep, and local T2), four model extraction methods (GIMME, iMAT, MBA, and mCADRE), and quantitative protection of metabolic functions (i.e., growth rate). First, we define a method to generate the ensemble of alternate solutions for each case. Next, we evaluate the growth rate predicted by all extracted context-specific models and determine that qualitatively protecting the biomass reaction (as previously suggested (Richelle et al., 2019a)) is not sufficient to accurately predict the experimentally measured growth rate. Following this, we quantify the variability in content of context-specific models in each ensemble in terms of conserved and variable pathways to assess the reproducibility of each method. Across all organisms and expression thresholds evaluated in this study, mCADRE generated the most reproducible models, whereas models generated by MBA showed the largest variance in reaction content. We also find that the size and content of models extracted using GIMME were the least sensitive to the applied expression threshold in all organisms evaluated in this study. We then demonstrate the utility of the receiver-operating-characteristic (ROC) plot in visualizing the performance of extracted context-specific models and propose a metric to select the model which best represents the biological system in the context of the application, using gene knockout data reserved from the model extraction dataset. Using a Euclidean distance metric, we quantified the proximity of the extracted models to the ideal model and found that GIMME generated the best-performing models for fast growing prokaryotes such as *E. coli*, whereas models extracted using mCADRE fared better in mammalian systems such as 786O. Finally, we establish a set of guidelines that an extracted model should satisfy for reliable hypothesis generation in biomedical and metabolic engineering applications.

## 2. RESULTS

### 2.1. Flux through required metabolic functions must be explicitly protected during model extraction

Model extraction methods aim to generate models that predict biologically relevant fluxes and accurately capture the sensitivity of the fluxome to genetic and environmental perturbations. Therefore, biologically relevant models must accurately recapitulate experimentally measured metabolite uptake and secretion rates and fluxes through required metabolic function (RMF) reactions. In this study, we consider the biomass formation reaction as an RMF reaction. Because the biomass reaction may not necessarily be retained in the extracted models, it should be protected as a core reaction to ensure retention (Richelle et al., 2019a). This was sufficient in optimization-based methods (GIMME and iMAT), in which fluxes were protected using lower and upper bounds in the metabolic model. However, protecting the biomass reaction was insufficient to ensure a biologically relevant growth rate in models extracted using MBA and mCADRE (Figure 1). Only 34 MBA models for *E. coli* generated using the 80^th^ percentile expression threshold predicted a growth rate greater than 90% of the experimentally measured growth rate (Supplementary Figure S1A). For 786O, only 36 of 500 models generated using MBA supported a growth rate within 10% of the maximum rate predicted by Recon2.2 (Supplementary Figure S1B). For CHO-S, only 9 of 500 generated MBA models predicted a growth rate within 10% of the maximum growth rate predicted by *i*CHO1766 (Supplementary Figure S1C). No model extracted using mCADRE for any organism correctly predicted biologically relevant growth rates despite protecting the biomass formation reaction itself as a core reaction. Core reactions in MBA and mCADRE are considered active if they can carry a flux of at least 10^−4^ mmol/gDW-h for *E. coli* or 10^−4^ mmol/gDW-day for 786O and CHO-S, which is several orders of magnitude less than the experimentally measured growth rate of all three organisms.

**Figure 1:**
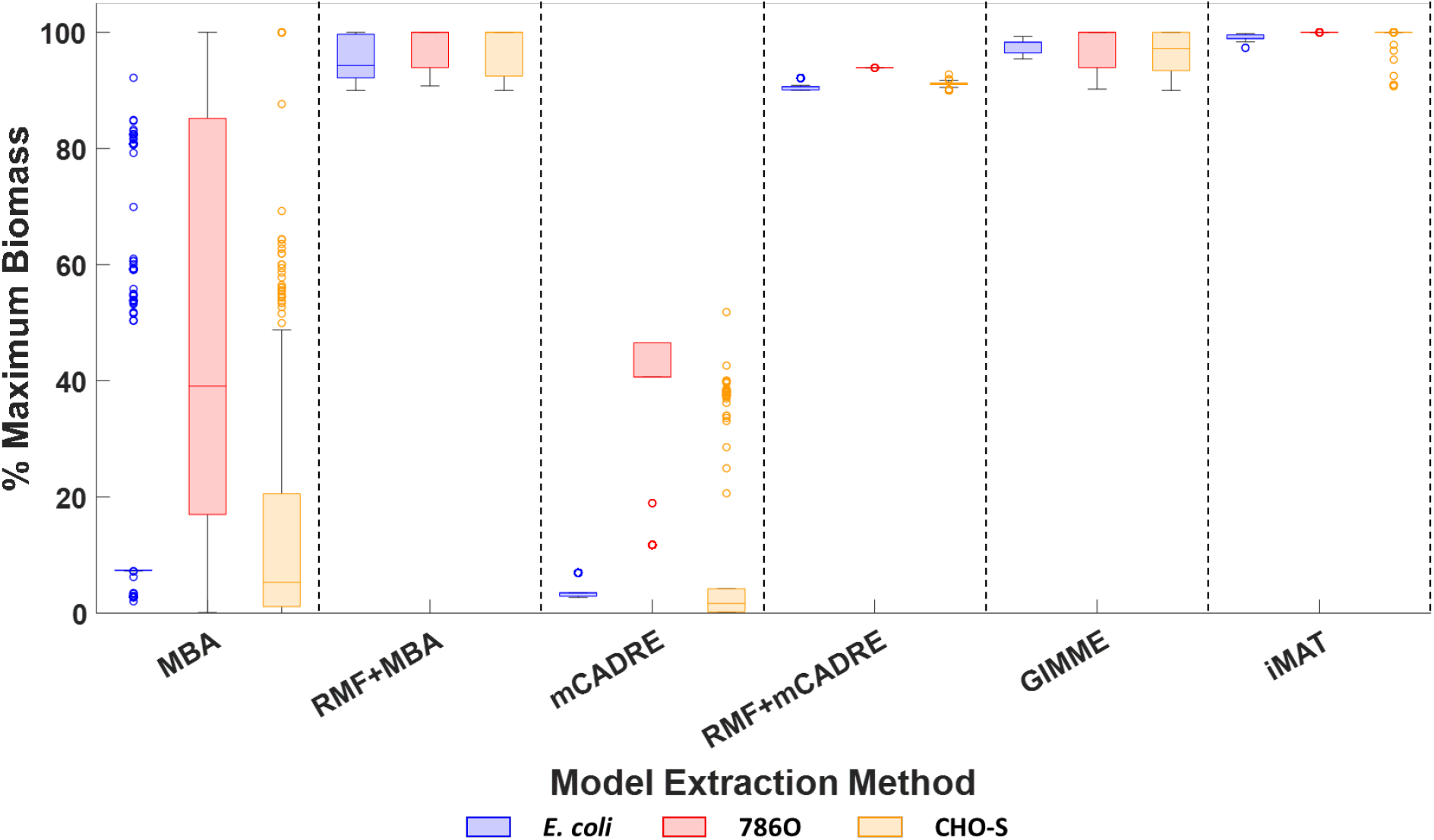
Retention of required metabolic functions. Box and Whisker plots show the distribution of the maximum growth rate predicted by extracted models relative to the maximum growth rate predicted by the genome-scale model for E. coli, 786O, and CHO-S using GIMME, iMAT, MBA, and mCADRE.

In *E. coli*, reactions from the electron transport chain (complexes I, II and III) and succinate dehydrogenase from the TCA cycle were necessary for ATP production but were inactivated because the associated transcript abundances were below the cutoff threshold. The resulting models therefore relied on the lower-yield substrate-level phosphorylation reactions for ATP generation and yielded lower growth rates compared to *i*JO1366. In 786O and CHO-S, reactions supporting cysteine and lysine uptake were removed based on transcriptomic evidence. Thus, the resulting models relied on *de novo* cysteine biosynthesis pathways and biocytin catabolism to meet the biosynthetic cysteine and lysine demands. The low abundance of biocytin in cell culture media limited lysine availability for protein synthesis, resulting in a considerably lower growth rate prediction compared to the respective parent genome-scale models. Ranking of non-core reactions based on expression scores prior to model pruning in mCADRE ensured that reactions required to sustain an experimentally measured growth rate were always removed due to low or missing gene expression values. However, very few MBA models fortuitously retained these reactions because MBA randomizes the removal order for reactions with low expression scores. Upon enforcing a mandatory minimum flux of 90% of the maximum growth rate predicted by the parent genome-scale model as a pruning criterion, all models generated by MBA and mCADRE predicted a biologically relevant growth rate for each of *E. coli*, 786O, and CHO-S (Figure 1). These findings suggest that even the most lenient threshold approaches such as StanDep and the Local T2 threshold can filter out reactions necessary to support key phenotypes and therefore, flux through RMF reactions must be explicitly protected during model extraction.

### 2.2. Choice of extraction method determines the scope of alternate solutions

Analysis of model sizes in each ensemble provided insights into the reproducibility and internal variability of model extraction methods. The ensemble generated using mCADRE showed the least dispersion in model sizes (average range = 2 for *E. coli*, 10 for 786O, and 14 for CHO-S), while models generated using MBA showed the largest dispersion in model sizes for *E. coli* (average range = 37) and CHO-S (average range = 280) (Figure 2, Supplementary Tables ST4, ST5, and ST6). For 786O, models generated using iMAT showed the largest size dispersion (average range = 128). Upon increasing the global expression threshold from the 60^th^ percentile to the 80^th^ percentile, the dispersion of model sizes from iMAT and MBA increased by up to 50%. However, ensembles generated using iMAT and MBA with StanDep or local T2 thresholding had lower size dispersion compared to models using global thresholding. The size dispersion correlated with the size of the core reaction set. For larger core reaction sets, model extraction methods choose pathways from a smaller set of non-core reactions for gap-filling, resulting in ensembles with smaller dispersions for thresholds with more core reactions. Interestingly, model size dispersion in ensembles generated using GIMME remained relatively unchanged in response to changes in threshold. On the other hand, rank-ordering of non-core reactions by mCADRE limits variability in removal order, and therefore, generated ensembles with the smallest size dispersion.

**Figure 2:**
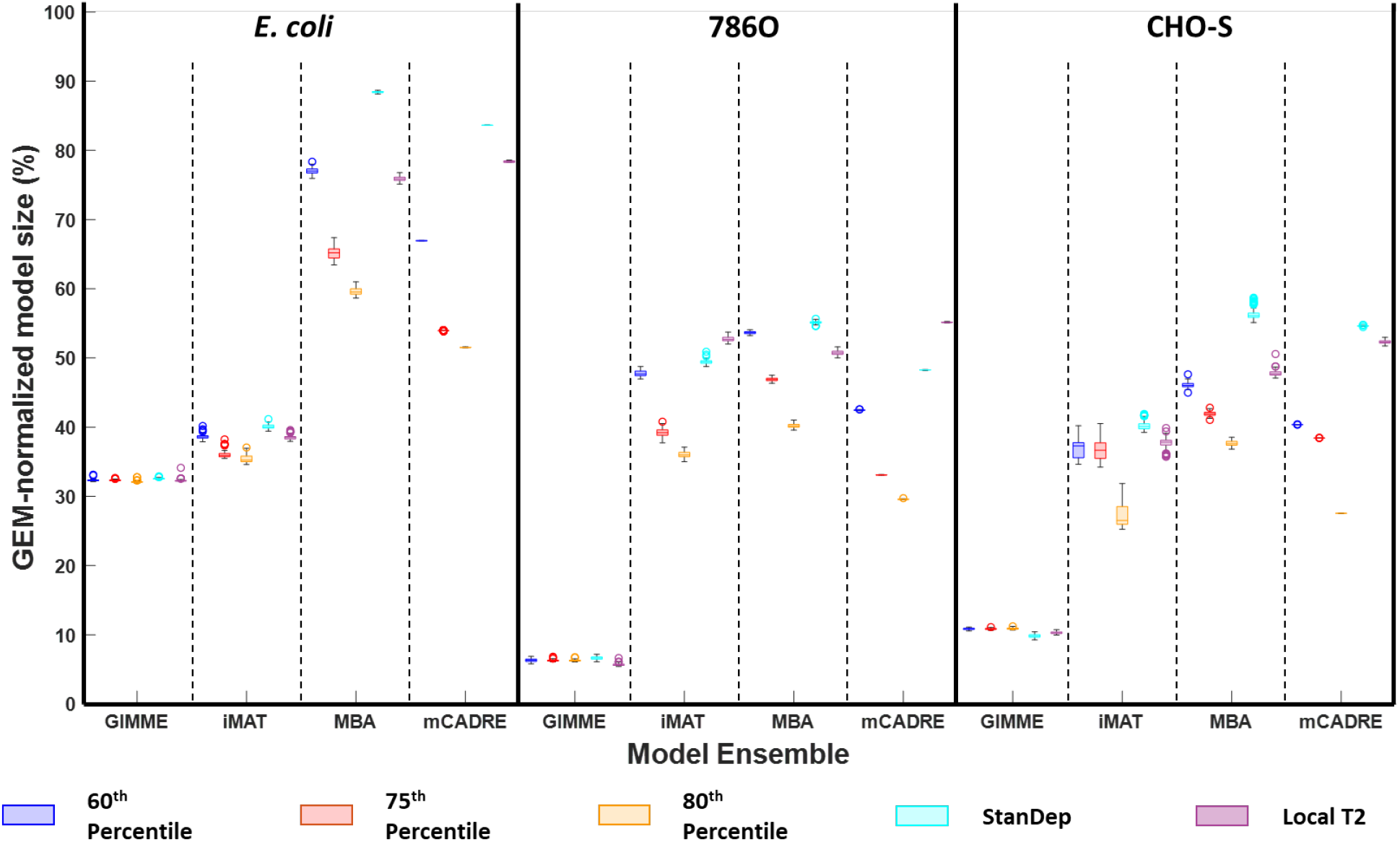
Size distribution of models in the ensemble generated using GIMME, iMAT, MBA, and mCADRE for E. coli, 786O and CHO-S with the global 60^th^ percentile threshold, global 75^th^ percentile threshold, global 80^th^ percentile threshold, StanDep, and the local T2 threshold.

Because a low size dispersion within an ensemble does not necessarily imply fewer alternate solutions, conserved and variable reactions in the ensemble must be identified and analyzed. During model extraction, we classified all reactions in the parent genome-scale models into one of four classes: conserved reactions (always retained in the ensemble), inactivated reactions (always removed in all models), variable reactions (retained in some models when certain criteria are met), and no data reaction (reactions lacking data in favor of retention or removal). The Jaccard index highlights the prevalence of each of these reaction classes and therefore quantifies the diversity of models within an ensemble.

The average Jaccard index for ensembles from mCADRE were 0.99, 0.99, and 0.98, in *E. coli*, 786O, and CHO-S, respectively. Over 98% of reactions in the extracted models were conserved reactions (Figure 3A). Upon varying the applied threshold, the number of conserved reactions in *E. coli* ranged from 872 to 1,426 reactions. The corresponding ranges were 1,722 to 3,199 reactions in 786O, and 1,161 to 2,249 reactions in CHO-S. Reactions were conserved in an ensemble because they were either core reactions, stoichiometrically coupled to core reactions, or stoichiometrically coupled to the biomass formation reaction. 434, 286, and 332 growth-coupled reactions were conserved in *E. coli*, 786O, and CHO-S, respectively. While only 315 reactions in *E. coli* were retained to activate blocked core reactions, this number increased up to 541 reactions in CHO-S and 1,019 reactions in 786O. This suggests that reaction retention in *E. coli* was primarily driven by biomass coupling, whereas gene expression data were the primary cause of reaction retention in the eukaryotic models. 27 reactions in *E. coli*, 303 reactions in 786O, and 259 reactions in CHO-S constituted alternate solutions (Figure 3B). In *E. coli*, these 27 reactions (21 reactions from glycerophospholipid metabolism, 3 metabolite transport reactions, and 3 reactions from lipopolysaccharide biosynthesis) were included to ensure flux consistency of seven core reactions (five transport reactions, and one reaction each from lipopolysaccharide and glycerophospholipid biosynthesis). In 786O, alternate solutions resulted from variability in 203 transport reactions, 34 glycosylation reactions, 22 reactions from fatty acid metabolism, and 8 reactions from nucleotide metabolism, 10 reactions from amino acid metabolism, and 23 reactions from central metabolism. These reactions were retained in the extracted models to activate 195 core reactions, primarily from fatty acid metabolism, all of which have four alternate pathways on average activating them. In CHO-S, 187 transport reactions, 25 reactions from fatty acid metabolism, 15 glycosylation reactions, 11 reactions from nucleotide metabolism, and 21 reactions from central and amino acid metabolism make up all identified alternate solutions. Similar to 786O, the core reactions activated by these non-conserved reactions are predominantly from fatty acid metabolism. Since mCADRE attempts to remove all non-core reactions, none of the reactions in the model were classified as no data reactions.

**Figure 3A:**
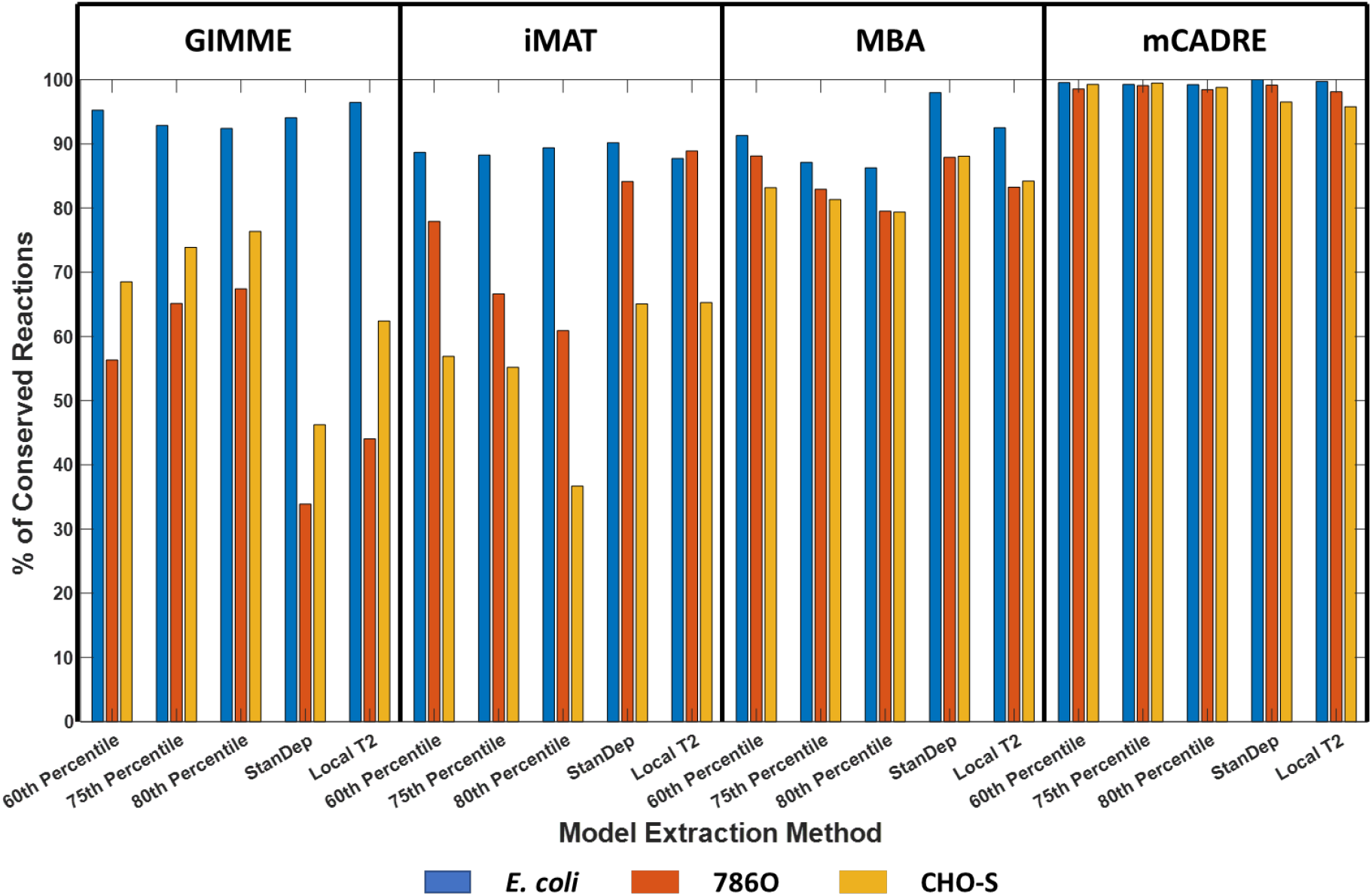
Fraction of conserved reactions in models extracted using GIMME, iMAT, MBA, and mCADRE for *E. coli*, 786O, and CHO-S with various thresholds.

**Figure 3B:**
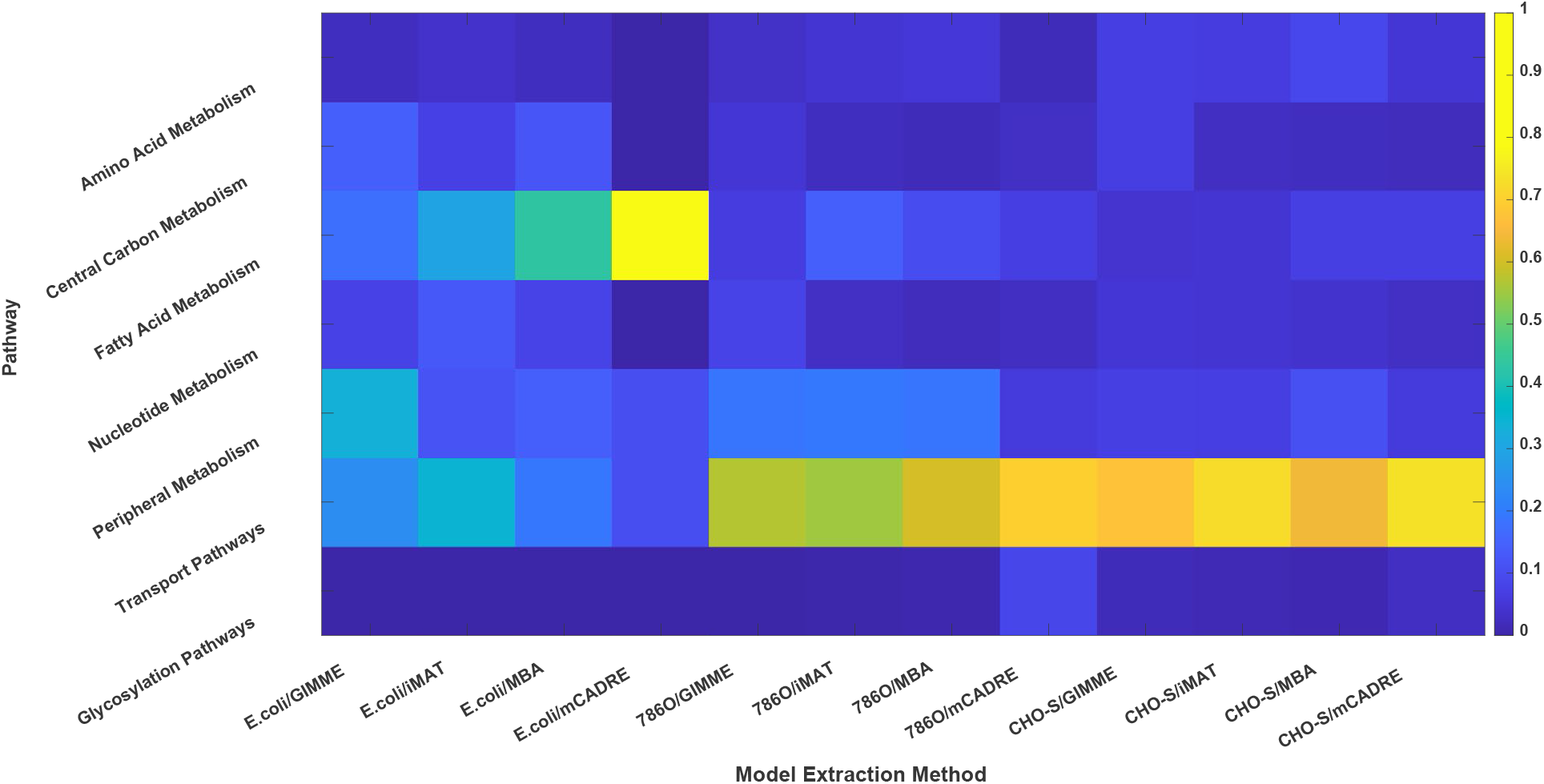
Fraction of reactions from various pathways (0 representing no variable reactions and 1 representing all variable reactions) contributing to alternate solutions in models extracted using GIMME, iMAT, MBA, and mCADRE for *E. coli*, 786O, and CHO-S with various thresholds

Compared to mCADRE, MBA ensembles had greater size dispersion and lower Jaccard index values (averaging 0.95 in *E. coli*, 0.86 in 786O, and 0.82 in CHO-S). Although MBA used more core reactions than mCADRE, an average 10% reduction in conserved reactions was observed in all three organisms. Unlike mCADRE, MBA permits removing core reactions if at least twice as many non-core reactions are removed. In addition, conserved reactions accounted for only 91%, 84%, and 83% of the extracted models for *E. coli*, 786O, and CHO-S, respectively. This contrasted with mCADRE, in which >99% of the reactions in all extracted models were conserved. The variable fraction of the models was considerably higher in MBA models compared to mCADRE models (Figure 4A), accounting for 247 reactions in *E. coli*, 1,436 reactions in 786O, and 1,579 in CHO-S, of which, 23 reactions in *E. coli*, 49 reactions in 786O, and 91 reactions in CHO-S were rendered growth-coupled by mCADRE. The variable reactions in extracted models were predominantly from fatty acid metabolism in *E. coli* and from metabolite transport pathways in 786O and CHO-S (Figure 3B). Of these variable reactions, 171 reactions in *E. coli*, 1,114 reactions in 786O, and 1,222 reactions in CHO-S were always removed in ensembles generated using mCADRE. This is because MBA randomizes the removal order of non-core reactions whereas mCADRE sorts non-core reactions based on expression and connectivity evidence prior to removal. Thus, certain non-core reactions are always eliminated by mCADRE because their low gene expression increases their removal priority, while MBA may retain them if competing non-core reactions are removed earlier. This implementation difference contributed to the larger variation in size and content in models extracted using MBA compared to other methods.

**Figure 4A:**
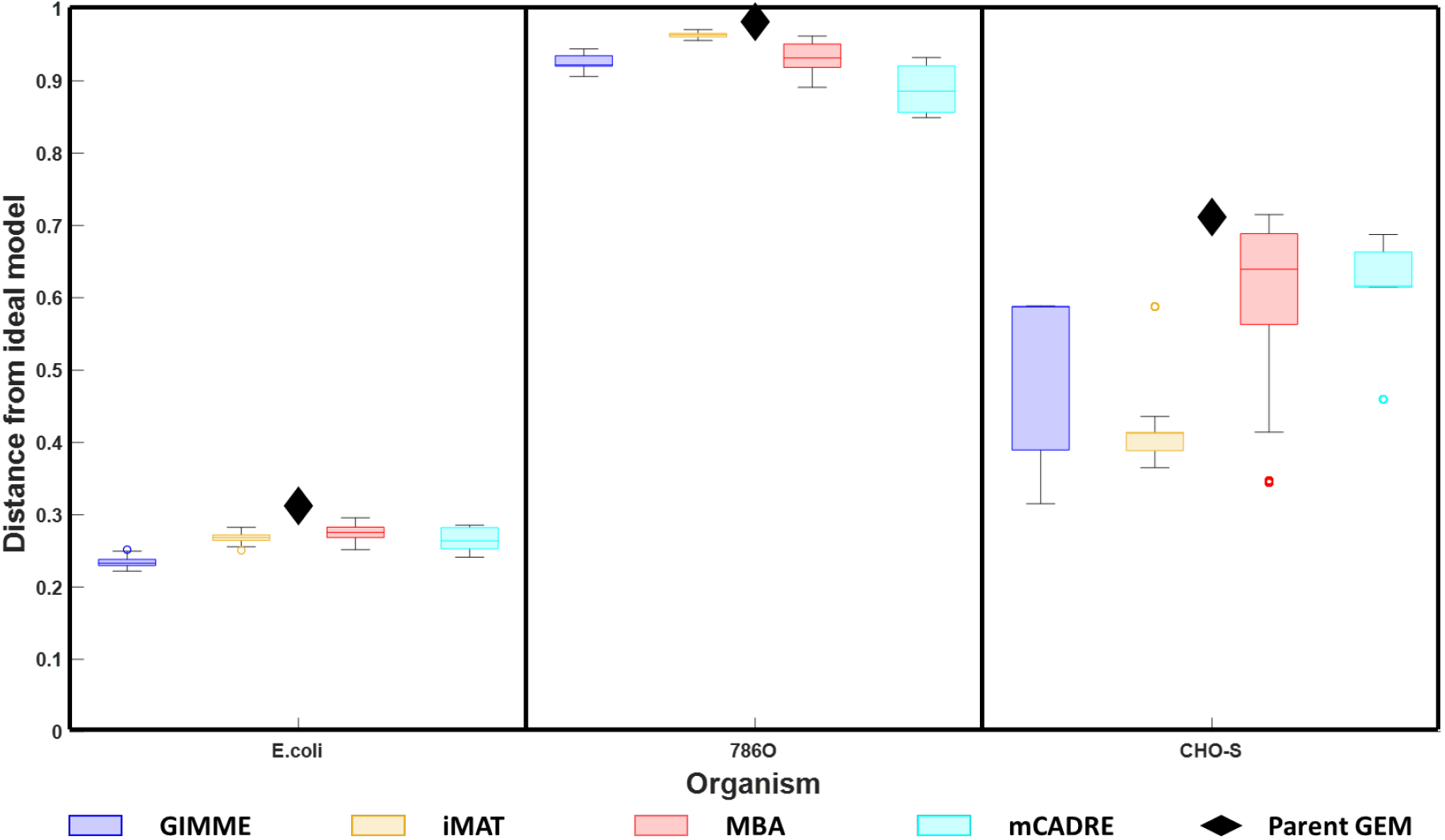
Improvement in quality of models extracted using GIMME, iMAT, MBA, and mCADRE for *E. coli*, 786O, and CHO-S compared to the parent genome-scale models. The ideal model correctly classifies all essential and non-essential reactions and therefore, has a specificity and sensitivity equal to 1. The distance from the ideal model is calculated as 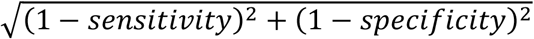.

**Figure 4B:**
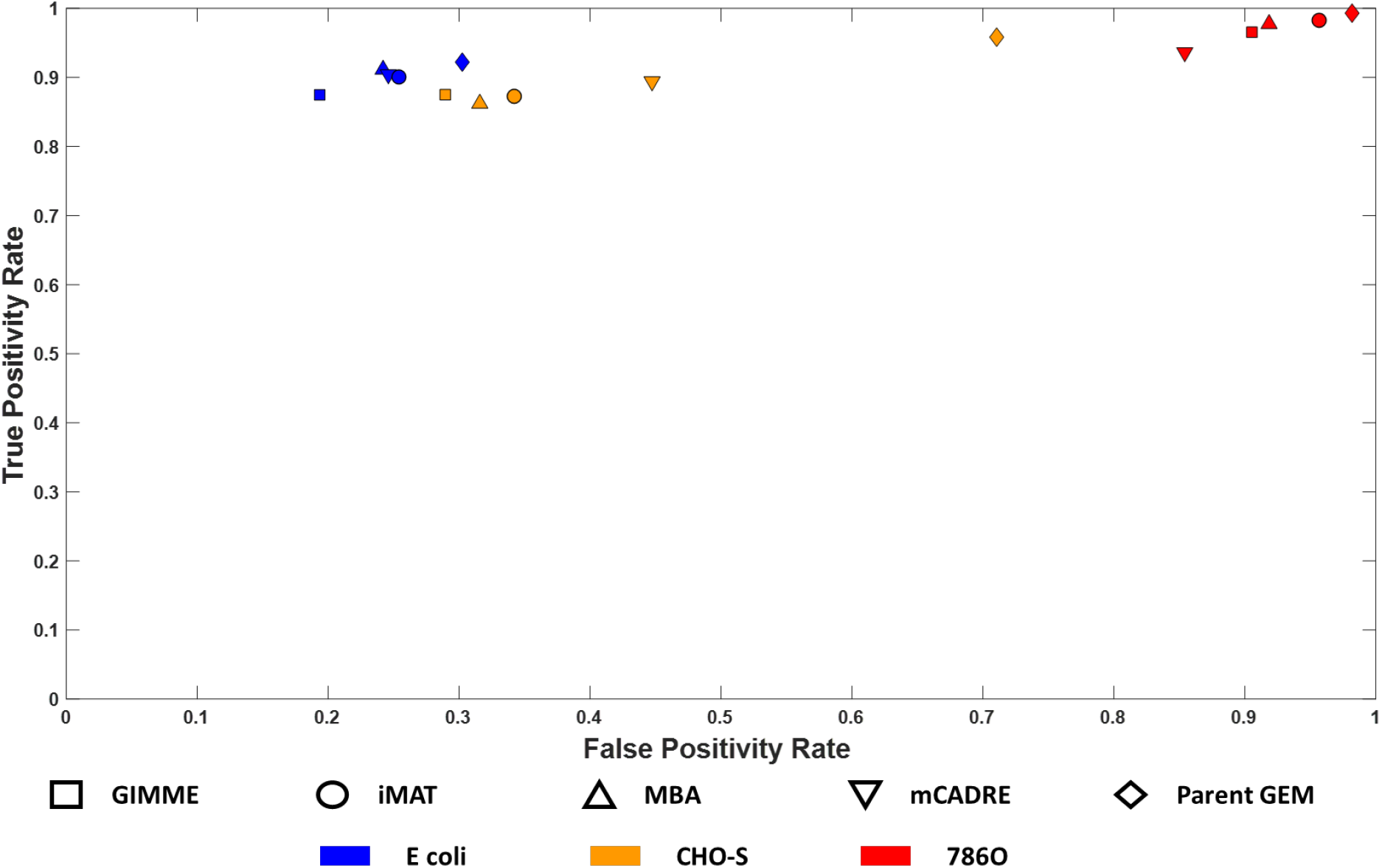
Receiver Operating Characteristic (ROC) plot showing the improvement in model performance of the best models extracted using GIMME, iMAT, MBA, and mCADRE relative to the parent genome-scale model in *E. coli*, 786O, and CHO-S.

Compared to MBA, iMAT models had fewer reactions, lower dispersion, and lower variability in model content with a Jaccard index of 0.96, 0.86, and 0.8 in *E. coli*, 786O, and CHO-S, respectively. Ensembles generated using iMAT for *E. coli* had the smallest fraction of conserved reactions (88%). For 786O and CHO-S, this fraction was 74% and 55%, respectively, considerably lower than mCADRE despite having the same number of core reactions. Unlike mCADRE, iMAT does not remove all reactions below the high expression threshold but attempts to inactivate only those reactions whose expression score is below the specified lower threshold. Moreover, iMAT permits removing core reactions if an equal number of low expression reactions were inactivated. Reactions from transport pathways and fatty acid metabolism accounted for 65% of all variable reactions in the *E. coli* ensembles (Figure 4B). Meanwhile, reactions from fatty acid metabolism, cofactor biosynthesis, and transport pathways accounted for 88% of the variable reactions in 786O, whereas reactions from metabolite transport pathways alone accounted for 70% of the variable reactions in CHO-S.

Although the GIMME ensembles had low size dispersions relative to iMAT and MBA, a pairwise comparison of models based on reaction content revealed that the scope of alternate solutions varied based on the topological features of the parent GSM model. Ensembles extracted using GIMME for *E. coli* had an average Jaccard index of 0.99 with 426 conserved reactions across the ensemble, 1,815 reactions always removed in all models, and 342 reactions contributing to alternate solutions. Of the 426 conserved reactions, 375 reactions were growth-coupled in *i*JO1366, 43 reactions were growth-coupled in the extracted models but not in *i*JO1366, one reaction (ATP maintenance) was retained based on pre-specified flux bounds, and six reactions from central metabolism were retained as alternatives to low-expression reactions. Of the 342 variable reactions, 224 reactions from metabolite transport, fatty acid metabolism, tryptophan biosynthesis and nucleotide phosphorylation pathways were growth-coupled when retained in the extracted models. Ensembles for both eukaryotic models had more diverse alternate solutions with an average Jaccard index of 0.72 for CHO-S and 0.64 for 786O. The number of conserved reactions was also reduced to 170 reactions in CHO-S and 83 reactions in 786O with only 127 and 44 reactions coupled to biomass formation in iCHO1766 and Recon2.2, respectively. 4,757 reactions in CHO-S and 5,861 reactions in 786O were inactivated in every extracted model. However, the number of variable reactions in each case increased to 1,736 reactions in CHO-S and 1,841 reactions in 786O, which is much greater than *E. coli*, despite similarities in model sizes in all three ensembles. 70% of these variable reactions were inter-compartment metabolite transport reactions, 10% from amino acid metabolism, 6% from fatty acid metabolism, and the remaining from cofactor biosynthesis and nucleotide biosynthesis and salvage. The primary objective of GIMME is to inactivate reactions with genes expressed below the threshold while ensuring that RMF reactions are retained and fully operational. Thus, we classify reactions as: (i) growth-coupled, (ii) low-expression, and (iii) maybe-on. All growth-coupled reactions are always retained in every extracted model. Low-expression reactions are always removed unless coupled to the RMF reaction. The inactivation of low-expression reactions forces flux through alternate pathways, when available, to meet the demands of the RMF reaction. Pathways that are the sole alternatives to low-expression reactions are retained in every extracted model. However, when alternate pathways exist, variable reactions can be retained, resulting in alternate solutions. Reactions with no available data have no reason for retention or removal and therefore contribute to alternate pathways. As such, alternate solutions from GIMME are determined predominantly by the topological features of the parent GSM. In *E. coli*, a much larger fraction of metabolism is growth-coupled leading to less diverse alternate solutions. However, models relying on more complex media, such as 786O and CHO-S have a more diverse set of alternate solutions.

### 2.3. ROC plots help evaluate the quality of extracted models

Diverse ensembles of context-specific models can be generated, but it is often unclear which models are most biologically relevant. To validate extracted models, gene dispensability data, flux redirections, and fluxomics datasets can be used (Opdam et al., 2017). Here we rely on gene knockout data to evaluate the quality of alternate optimal models. The ideal model would correctly identify all essential and non-essential genes. Integrating transcriptomics data deactivates pathways that are inactive in the context of interest and is therefore expected to reconcile false predictions by the genome-scale model. Here we evaluate the specificity and sensitivity using receiver operating characteristic (ROC) plots (see Methods section for the definition of specificity and sensitivity and Supplementary Figure S2 ROC plots for *E. coli*, 786O, and CHO-S). After computing the specificity and sensitivity for each model, the distance from the ideal model was computed and then compared with the parent genome-scale model.

All extracted models outperformed their respective parent GEM models in predicting gene dispensability. This is because model pruning removes alternate routes that compensate for the loss of function of essential reactions, which reconciles false-positive predictions in the genome-scale model. We find that GIMME models had the highest specificity for *E. coli* and CHO-S with an average sensitivity of 0.87 and 0.71, respectively. mCADRE generated the highest specificity models for 786O with an average specificity of 0.14. The best models generated for *E. coli* and CHO-S using GIMME showed a 29% and 55% improvement in gene essentiality predictions compared to *i*JO1366 and *i*CHO1766, respectively. On the other hand, the best model for 786O generated using mCADRE only showed a 13% improvement compared to Recon2.2.

The essentiality of 203 genes were reconciled in the best performing model generated using GIMME for *E. coli*, including 30 genes from fatty acid biosynthesis, nucleotide biosynthesis, and glycolysis. Compared to other models in the ensemble, the best performing model failed to reconcile the essentiality of the b1638 gene that encodes the PDX5POi reaction involved in pyridoxal phosphate biosynthesis. The PDX5PO2 reactions serves as an alternate route to pyridoxal phosphate synthesis when the PDX5POi gene is inactivated. Because PDX5PO2 is not associated with any gene, it is not preferentially removed or retained in models generated using GIMME and iMAT, due to which, b1638 is always reconciled in these ensembles. In contrast, PDX5PO2 is treated as a low confidence reaction by MBA and mCADRE, leading to prioritized removal. As a result of this, MBA and mCADRE can reconcile the essentiality of b1638.

The essentiality of 62 genes predominantly from fatty acid metabolism and transport pathways were reconciled in the best performing model for 786O generated using mCADRE. In the best model for CHO-S constructed using GIMME, the essentiality of 18 genes from fatty acid metabolism and the TCA cycle were reconciled. The best models generated for 786O and CHO-S reconciled all essential genes reconciled in their respective ensembles.

The difference in gene essentiality reconciliation between the three models is attributable to differences in the metabolism of *E coli* and mammalian cells, which are reflected in the topological features of *i*JO1366, Recon2.2, and iCHO1766. Because *E. coli* grows in minimal media, a large fraction of its metabolism is biosynthetic, leading to a higher number of growth-coupled pathways. Protection of flux through the biomass reaction leads to removal of only dispensable pathways supported by low gene expression in models extracted using GIMME. This gave rise to models with the largest increase in specificity compared to the parent genome-scale model in *E. coli*. On the other hand, because a much smaller fraction of Recon2.2 and iCHO1766 is coupled to biomass production, removal of reactions without evidence-based prioritization leads to erroneous removal of essential reactions. This resulted in models with low specificity in 786O and CHO-S. In contrast, mCADRE prioritizes removal of reactions that are poorly expressed and weakly connected to highly expressed reactions. This systematic removal protects against the removal of highly expressed reactions in potentially essential pathways, thereby generating models with higher specificity than those extracted using GIMME for 786O. In comparison, models generated by iMAT and MBA did not perform as well as those generated by GIMME as suggested by their proximity to the parent genome-scale model (Figure 4 and Supplementary Figure S2). Models generated by iMAT were much closer to the parent genome-scale model for *E. coli* and 786O, but performed considerably better in CHO-S.

## 3. DISCUSSION

This study evaluates key parameters influencing the quality of context-specific models extracted with various methods using gene expression data. While the choice of model extraction method and the threshold for gene expression remain the most important factors affecting model size, our analysis reveals that depending on the choice of model extraction method, the exploration of alternate solutions can lead to drastically different models. These findings suggest the need for a set of guidelines for extracting the most meaningful and biologically relevant context-specific models, to supplement guidelines on model construction (Thiele and Palsson, 2010), model annotation (Ebrahim et al., 2015), and model parameterization (Schinn et al., 2021b). Key guidelines are presented in Table 1, a workflow incorporating the proposed guidelines is shown in Figure 5, and the steps to implement the workflow are listed in Table 2. Three steps (Figure 5) are involved in the extraction of context-specific models from genome-scale models: (i) pre-processing, (ii) ensemble generation, and (iii) ensemble screening. The pre-processing step transforms the raw model and transcriptomic data into a format compatible with model extraction methods.

**Table 1:**
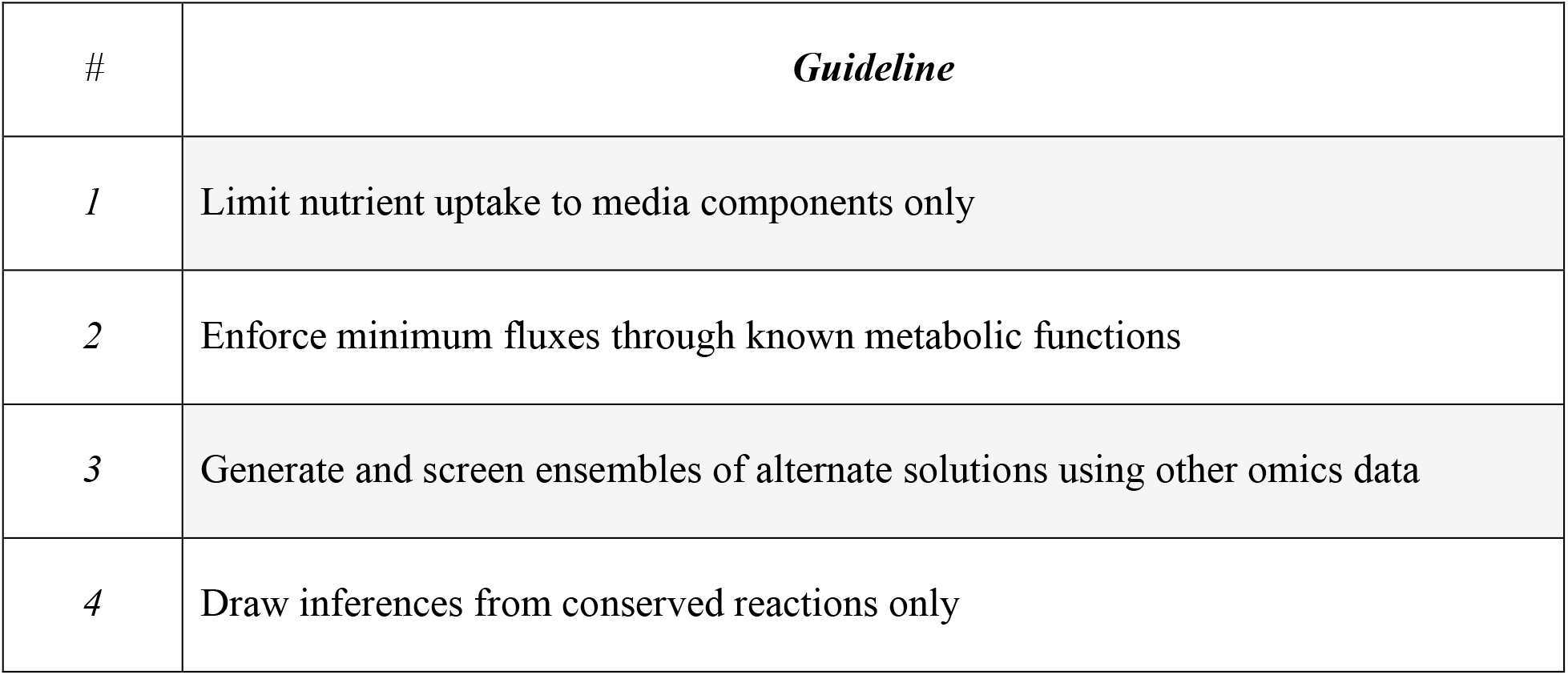
Guidelines for extracting meaningful metabolic models using transcriptomics data

**Table 2:** Implementation of the workflow depicted in Figure 5.

| STEP | DESCRIPTION |
| --- | --- |
|  | **Model preprocessing** |
| **STEP 1A:** | Impose the lower and upper bounds for the uptake and secretion of all measured metabolites as well as the growth rate. For metabolites in the growth medium that are not measured, an arbitrary bound limiting their uptake can be imposed. Identify all reactions incapable of carrying flux using Flux Variability Analysis and remove them. The resultant pre-processed model should be flux consistent. |
|  | **Data preprocessing** |
| **STEP 1B:** | Compute reaction expression scores from gene expression data using defined reaction-specific Gene-Protein-Reaction (GPR) rules.<br>Generate multiple core reaction sets by applying different thresholds to the computed reaction expression scores. Local thresholding methods are often preferred due to their ability to retain lowly expressed housekeeping genes. |
|  | **Identify metabolic tasks that define the cell's phenotype** |
| **STEP 2:** | Generate a list of metabolic tasks that must be retained in extracted models. Metabolic tasks with available experimental measurements must be quantitatively protected. Other identified metabolic tasks should be added to the sets of core reactions. |
|  | **Generate ensembles of context-specific models** |
| **STEP 3:** | Using the preprocessed model from step 1a, the preprocessed reaction expression scores from step 1b, and the metabolic tasks from step 2 as inputs, generate ensembles of at least 50 models using any model extraction method. |
|  | **Screen and select the best-performing models** |
| **STEP 4:** | For each model in the generated ensemble, compute the specificity and sensitivity using validation data (gene knockout, flux prediction, etc.). Compute the distance from the ideal model using the expression: $\sqrt{(1 - sensitivity)^2 + (1 - specificity)^2}$. The top performing models have the lowest distance metric. |

**Figure 5:**
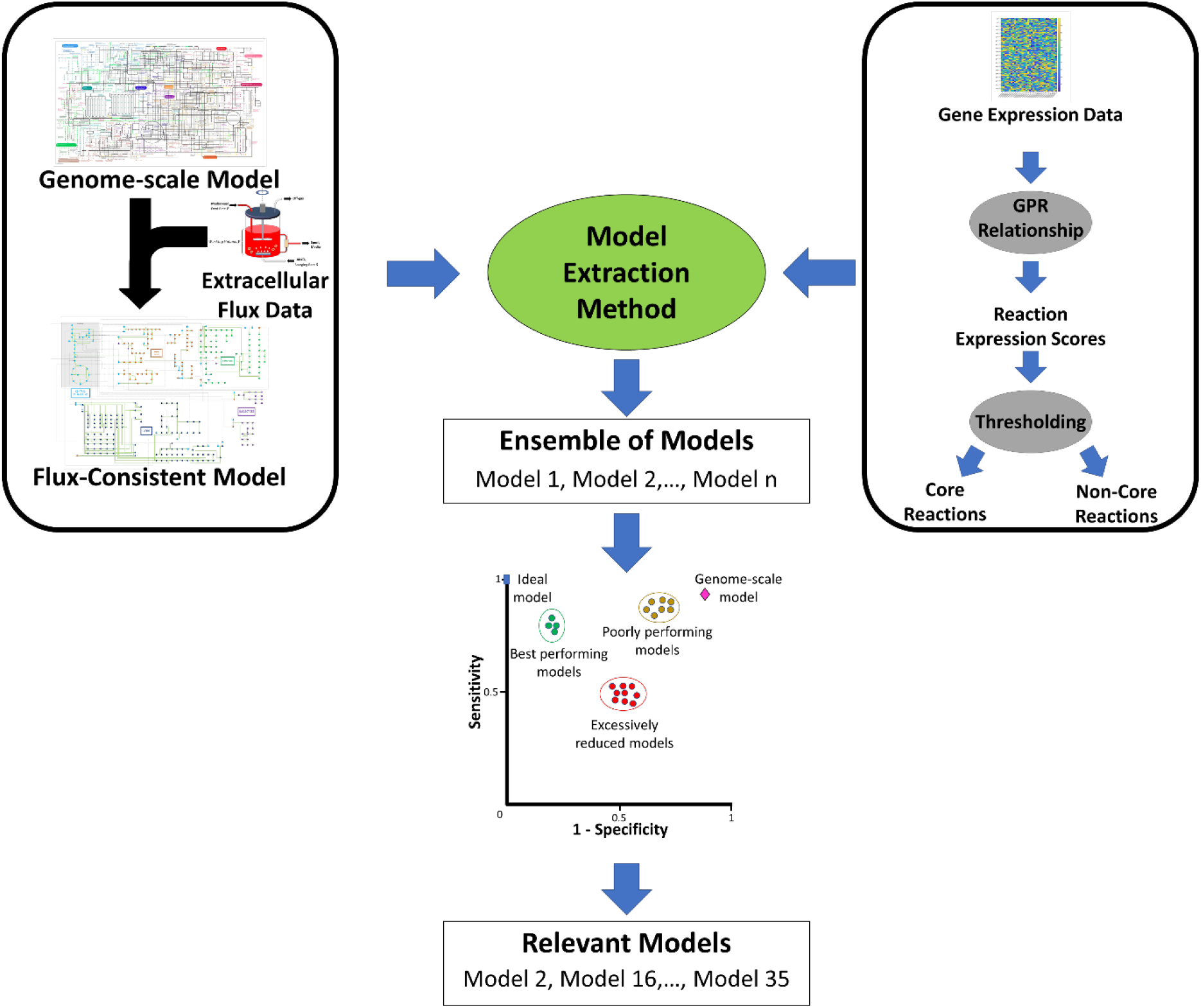
Generalized workflow pipeline for extracting context-specific models using gene-expression data

Preprocessing of transcriptomics involves applying a threshold to determine which reactions are likely active. To this end, transcriptomic data are log-transformed and mapped to reactions via gene-protein-reaction (GPR) relationships. A threshold (top 25^th^ percentile, top 50^th^ percentile, etc.) is applied to reaction expression scores to extract lists of reactions based on the requirements of model extraction methods. Here we investigated combinations of five thresholds (global 60^th^ percentile, global 75^th^ percentile, global 80^th^ percentile, StanDep, and local T2 threshold) and four model extraction methods (GIMME, iMAT, MBA, and mCADRE). GIMME and mCADRE require the lists of reactions with expression scores below and above the specified threshold, respectively. iMAT and MBA require two thresholds to classify reactions into highly expressed and weakly expressed sets. Incorporating media information identifies and eliminates inconsistent core reactions which protects the workflow from extraction failures (see Supplementary Results). After preprocessing, gap-filling of metabolic networks is performed using model extraction methods to ensure flux consistency of the core reaction set.

During model extraction, it is mandatory to retain and protect the flux through known metabolic functions in the conditions being investigated. Indeed, required metabolic functions are not always retained in extracted models (Opdam et al., 2017) and protecting metabolic functions reduces the variability in model content between models extracted using different extraction methods (Richelle et al., 2019a). This study, however, finds that merely protecting these tasks is insufficient to ensure the required flux through the metabolic task. For example, the predicted growth rate in *E. coli* drops by over 99% in models generated using mCADRE when a minimum growth rate is not enforced. This suggests that while gene expression data provides insights into pathway activity, it alone is insufficient to distinguish between the various metabolic states underpinning the metabolic task. Although a comprehensive list of condition-specific metabolic tasks may be obtained through a literature search, sets of metabolic known tasks in rat and human tissues have been published (Blais et al., 2017; Richelle et al., 2019b; Thiele et al., 2013). Furthermore, context-specific metabolic tasks can be predicted from transcriptomic data to inform which of all tasks should be protected when extracting a model for the desired conditions or cell type (Masson et al., 2022; Richelle et al., 2019a; Richelle et al., 2021). The inability to consistently retain and predict a required flux through essential metabolic functions implies that flux constraints on these reactions complement gene expression data and improve the biological relevance of extracted models.

The size, content, and predictive capabilities of the model are strongly influenced by the choice of model extraction method and the applied threshold for gene expression, as seen in previous studies (Opdam et al., 2017; Richelle et al., 2019b). Therefore, the choice of the right combination of parameters is crucial for extracting a meaningful model. Here we demonstrated that ROC plots can be used to identify the best performing models. While models generated using individual gene-specific local thresholds (Uhlen et al., 2015) or thresholds derived from hierarchical clustering (Joshi et al., 2020) were generally better, these thresholding methods can only be applied when multiple gene expression data samples are available. In addition to gene knockout data used for screening in this study, other types of biological data such as metabolomics and fluxomics data can be used for validation so long as the model’s recapitulation of the validation dataset can be represented using a confusion matrix. While metabolomics data reveals which metabolites actively participate in the condition being investigated, fluxomics data elucidates pathway utilization to validate generated models. Furthermore, the quality of models extracted using different algorithms varied based on the biology of the organism in question. Using available gene knockout data, we found that GIMME generated the best performing models in fast-growing prokaryotes such as *E. coli*, whereas the corresponding models generated for a function-oriented cell such as 786O were sub-par. These differences suggest the need for a careful assessment of thresholds and methods while constructing context-specific models for targeted applications.

The impact of alternate solutions must be assessed while extracting and/or and developing tools to extract context-specific models. Alternate optima provide meaningful insights into the reproducibility of the algorithm and highlight the variable parts of the extracted metabolic networks (Rossell et al., 2013). This arises from the insufficiency of available gene expression data to resolve pathway usage in those parts of metabolism. Thus, any inferences drawn from flux distributions involving those pathways are potentially ambiguous and would require additional validation. Furthermore, for algorithms of lower reproducibility such as MBA, generation of an ensemble of models increases the likelihood of identifying better performing models that may be more relevant to the condition being investigated.

An important factor affecting the performance of extracted models is the quality of the parent genome-scale model. While curated models such as those for *E. coli* benefit from a wealth of available literature, thereby leading to models with very high specificity and sensitivity, less studied and more complex organisms do not enjoy the same luxury. For example, the parent genome-scale model for 786O, Recon2.2, has a very low sensitivity of 0.02. This indicates a need for developing algorithms that leverage gene knockout data in addition to gene expression data for extracting accurate context-specific models. Better model extraction algorithms that can accurately capture the biological state of the cell will simplify the model reduction step commonly performed before computationally intensive analyses such as 13C-MFA (Sacco and Young, 2021), kinetic modeling (Islam et al., 2021), hybrid models(Khaleghi et al., 2021), and models integrating other cell processes with metabolism, such as signaling pathways, protein secretion, and many other processes (Elsemman et al., 2022; Gutierrez et al., 2020; Karr et al., 2012). This will expand the coverage of biological data that can be integrated with metabolic models to gain novel insights into the biology of the organism, study the progression of diseases, identify novel therapeutics, and inform metabolic engineering strategies in production hosts.

## 4. Methods

### 4.1. Models and Data Sources

The metabolic models *i*JO1366 (Orth et al., 2011), Recon 2.2 (Swainston et al., 2016), and *i*CHO1766 (Hefzi et al., 2016) for *E. coli*, human metabolism, and Chinese hamster ovary (CHO-S) cells were used as parent genome-scale models for extraction of context-specific models. Published glucose uptake rate, growth rate, and acetate secretion rate for *E. coli* grown in M9 Minimal Medium were used (Leighty and Antoniewicz, 2013). Glucose uptake rate, lactate secretion rate, growth rate, and uptake and secretion rates for amino acids were obtained from the NCI-60 database for the 786O renal cancer cell line (Jain et al., 2012; Opdam et al., 2017) and from literature for the CHO-S cell line (Hefzi et al., 2016). Gene expression data for *E. coli* grown in M9 minimal medium, 786O, and CHO-S were obtained from previously published data by Monk et al. (2016), the NCI-60 database (Klijn et al., 2015), and previously published data by Hefzi et al. (2016), respectively.

### 4.2. Model and Data Preprocessing

Gene expression data were converted to reaction expression scores using a gene-protein-reaction (GPR) relationship. A GPR relationship is a Boolean expression that relates genes products to enzymes catalyzing a reaction. An OR relationship indicates that a reaction can be catalyzed by multiple isozymes. In this case, the reaction expression score is computed as the maximum expression of the genes encoding the different isozymes. Association of multiple subunits is modeled using the AND relationship. The reaction expression score for an AND relationship is evaluated as the minimum expression of the genes encoding the various subunits. Reactions without GPR relationships or with missing gene expression data were assigned an expression score of -1. These scores were used to identify global thresholding approaches. Expression scores using StanDep were computed as described by Joshi et al. (2020) whereas local T2 thresholding was performed as described by Richelle et al. (2019b). These approaches enable the better retention of more lowly expressed housekeeping genes and reactions (Joshi et al., 2022). Flux variability analysis (Mahadevan and Schilling, 2003) was performed to identify and remove inactive reactions so that all reactions in the parent models used for transcriptomics-based model extraction are flux consistent.

### 4.3. Model Extraction Methods

GIMME (Becker and Palsson, 2008) requires as inputs one expression threshold and assignment of a reaction as the required metabolic function (RMF). Values corresponding to the 60^th^, 75^th^, and 80^th^ percentile in the reaction expression scores were applied as thresholds to determine which reactions must be removed. For expression scores computed using StanDep and the local T2 approach, thresholds of 0 and 5*ln(2), respectively were applied. The biomass reaction was selected as the RMF reaction for all three organisms and a mandatory minimum of 90% of the maximum growth rate was enforced during model extraction. Since GIMME solves a linear programming problem to identify context-specific models, alternate solutions were identified by imposing an integer cut that eliminates previously identified solutions (Maranas and Zomorrodi, 2016).

iMAT (Zur et al., 2010) requires one threshold for high expression reactions and one for low expression reactions. For the global thresholding cases, expression scores corresponding to the 60^th^, 75^th^, and 80^th^ percentile were used to identify core reactions that must be included in the extracted model, whereas scores corresponding to the 20^th^ percentile were considered inactive reactions for removal. For StanDep and the local T2 cases, equal upper and lower threshold of 1 and 5*ln(2), respectively were applied. Because iMAT does not inherently protect flux through the RMF reaction, a lower bound of 90% of the maximum biomass flux was enforced in the MILP formulation of the iMAT case. As with GIMME, alternate solutions were identified using integer cuts.

MBA (Jerby et al., 2010) requires two sets of reactions be provided as inputs: one set corresponding to high confidence reactions that must be included in the extracted model and a medium confidence set that is maximally retained. For the global thresholding cases, reactions with scores above the 60^th^, 75^th^, and 80^th^ percentile were considered high confidence reactions whereas those with scores above the 40^th^ percentile but not part of the high confidence set were included in the medium confidence set. For StanDep, reactions with expression score greater than 110% of that method’s cluster threshold were considered high confidence reactions and reactions with expression scores between 90% and 110% were considered medium confidence reactions (Joshi et al., 2020). For the local T2 case, reactions with scores above the 75^th^ percentile were high confidence reactions and those with scores greater than 5*ln(2) and below the 75^th^ percentile were included in the medium confidence set. Alternate solutions were generated by permuting the removal order of low confidence reactions. In addition to ensuring flux consistency of the high expression reaction set, a minimum flux of 90% of the maximum growth rate was enforced as a criterion for removing reactions to ensure that all models in the ensemble can predict a biologically meaningful growth rate. A separate ensemble was also generated using the conventional implementation of MBA in which the biomass formation reaction is added to the set of high confidence reactions.

mCADRE (Wang et al., 2012) requires ubiquity scores to be provided as an input. Ubiquity scores for the global threshold cases were computed by normalizing reaction expression scores by the applied global threshold. Ubiquity scores for StanDep were computed as previously described by Joshi et al. For the local T2 case, ubiquity scores were calculated by normalizing expression scores to 5*ln(2) after applying appropriate local thresholds. Reactions with a ubiquity score greater than 1 were flagged as core reactions to be protected during model extraction. Because mCADRE ranks non-core reactions based on expression and connectivity evidence, only a subset of non-core reactions of equal rank can be permuted. Alternate solutions were identified by permuting the removal order of this subset of reactions. As with MBA, a minimum of 90% of the maximum growth rate was enforced as an additional criterion for model pruning. An ensemble was also generated using conventional mCADRE with the biomass formation reaction added to the set of core reactions.

All algorithms were implemented in the COBRA Toolbox (Heirendt et al., 2019) in MATLAB ^®^.

### 4.4. Analysis of Ensembles

The similarity of two models (*model*_*i*_ and *model*_*j*_) in any ensemble is quantified using the Jaccard Index defined as follows:

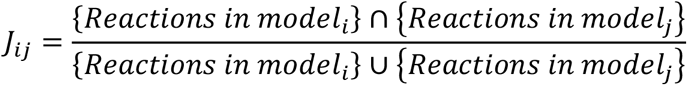

### 4.5. Validation of Extracted Models

Gene essentiality data inferred from gene knockout studies were used to screen ensembles of context-specific models. In silico gene essentiality was determined by computing the reduction in the growth rate upon inactivating one gene at a time in every extracted context-specific model. Genes were considered *in silico* essential if the predicted growth rate in the knockout model fell below 5% of the growth rate predicted by the original context-specific model. The quality of extracted context-specific models was evaluated by comparing model predictions of gene essentiality with experimentally determined gene essentiality. Gene essentiality data for WT *E. coli* grown in M9 Minimal medium was obtained from the KEIO collection (Baba et al., 2006). For the 786O cell line, gene essentiality was determined based on the CERES scores published in the NCI-60 database (Meyers et al., 2017). Genes with a CERES score less than zero were considered essential. The list of essential genes in CHO was obtained from (Xiong et al., 2021). Genes correctly predicted as non-essential were classified as true positive (TP) predictions, incorrectly predicted as essential were classified as false negative (FN) predictions, correctly predicted as essential were classified as true negative (TN) predictions, whereas those incorrectly predicted as non-essential were classified as false positive (FP) predictions. The specificity and sensitivity of the models were computed using the following expressions.

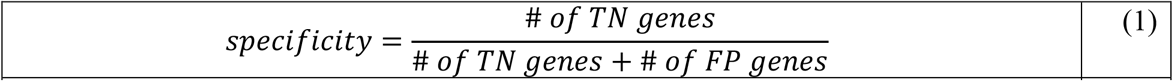

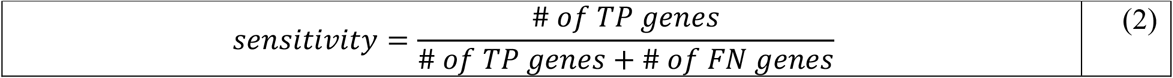

All extracted models and gene dispensability predictions are reported in the supplementary material.

## Acknowledgements

This work was supported by funding generously provided by Amgen, the Novo Nordisk Foundation (NNF20SA0066621) and NIGMS (R35 GM119850).

## Figure Captions

**Figure 1**

Retention of required metabolic functions. Box and Whisker plots show the distribution of the maximum growth rate predicted by extracted models relative to the maximum growth rate predicted by the genome-scale model for E. coli, 786O, and CHO-S using GIMME, iMAT, MBA, and mCADRE.

**Figure 2**

Size distribution of models in the ensemble generated using GIMME, iMAT, MBA, and mCADRE for E. coli, 786O and CHO-S with the global 60^th^ percentile threshold, global 75^th^ percentile threshold, global 80^th^ percentile threshold, StanDep, and the local T2 threshold.

**Figure 3**

(A) Fraction of conserved reactions in models extracted using GIMME, iMAT, MBA, and mCADRE for *E. coli*, 786O, and CHO-S with various thresholds.

(B) Fraction of reactions from various pathways (0 representing no variable reactions and 1 representing all variable reactions) contributing to alternate solutions in models extracted using GIMME, iMAT, MBA, and mCADRE for *E. coli*, 786O, and CHO-S with various thresholds

**Figure 4**

(A) Improvement in quality of models extracted using GIMME, iMAT, MBA, and mCADRE for *E. coli*, 786O, and CHO-S compared to the parent genome-scale models. The ideal model correctly classifies all essential and non-essential reactions and therefore, has a specificity and sensitivity equal to 1. The distance from the ideal model is calculated as 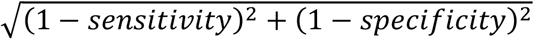.

(B) Receiver Operating Characteristic (ROC) plot showing the improvement in model performance of the best models extracted using GIMME, iMAT, MBA, and mCADRE relative to the parent genome-scale model in *E. coli*, 786O, and CHO-S.

**Figure 5**

Generalized workflow pipeline for extracting context-specific models using gene-expression data

