## Supplementary material for "Guidelines for extracting biologically relevant context-specific metabolic models using gene expression data": GIMME_multi.m

```
function [removed_reactions,gimme_objective,interm_model] = GIMME_multi(model, expressionRxns, threshold, obj_frac,N)
% Use the GIMME algorithm (`Becker and Palsson, 2008`) to extract a context
% specific model using data. GIMME minimizes usage of low-expression
% reactions while keeping the objective (e.g., biomass) above a certain
% value. Note that this algorithm does not favor the inclusion of reactions
% not related to the objective.
%
% USAGE:
%
%    tissueModel = GIMME(model, expressionRxns, threshold)
%
% INPUTS:
%
%    model:               input model (COBRA model structure)
%    expressionRxns:      expression data, corresponding to model.rxns (see
%                         mapGeneToRxn.m)
%    threshold:           expression threshold, reactions below this are
%                         minimized
%
% OPTIONAL INPUT:
%    obj_frac:            minimum fraction of the objective(s) of model
%                         (default value - 0.9)
%    N       :            Number of generated models (default = 1)
%                         Include N as an optional input if you want to
%                         generate an ensemble of N models consistent with
%                         the provided transcriptomics data. The first
%                         model is generated using the standard GIMME
%                         procedure. the remaining N-1 models are
%                         gnerated using integer cuts.
%
% OUTPUTS:
%    tissueModel:         extracted model
%
% `Becker and Palsson (2008). Context-specific metabolic networks are
% consistent with experiments. PLoS Comput. Biol. 4, e1000082.`
%
% .. Author: - Originally written by Becker and Palsson, adapted by S. Opdam and A. Richelle - May 2017
% .. Adapted by S. Gopalakrishnan - March 2020 - Included the option to generate alternate solutions

 f = waitbar(0,'Initializing GIMME runs');
figx = uifigure;
uT = uitable(figx);
uT.ColumnName = {'Model' 'Objective' 'Retained Reactions' 'Removed Reactions' 'Time (min)'};   
    if nargin < 4 || isempty(obj_frac)
        obj_frac = 0.9;
    end
    
    if nargin < 5
        N = 1;
    end
    

    objectiveCol = [find(model.c) obj_frac]; 

    nRxns = size(model.S,2);
    
    % Initialize Outputs:
    removed_reactions = zeros(nRxns,N);
    gimme_objective = zeros(N,1);
    
    %first make model irreversible
    [modelIrrev,matchRev,rev2irrev,irrev2rev] = convertToIrreversible(model);

    nbExpressionRxns = size(expressionRxns,1);
    if (nbExpressionRxns < nRxns)
        display('Warning: Fewer expression data inputs than reactions');
        expressionRxns(nbExpressionRxns+1:nRxns,:) = zeros(nRxns-nbExpressionRxns, size(expressionRxns,2));
    end

    nIrrevRxns = size(irrev2rev,1);
    expressionRxnsIrrev = zeros(nIrrevRxns,1);
    for i=1:nIrrevRxns
        expressionRxnsIrrev(i,1) = expressionRxns(irrev2rev(i,1),1);
    end

    nObjectives = size(objectiveCol,1);
    for i=1:nObjectives
        objectiveColIrrev(i,:) = [rev2irrev{objectiveCol(i,1),1}(1,1) objectiveCol(i,2)];
    end

    %Solve initially to get max for each objective
    for i=1:size(objectiveCol)
        %define parameters for initial solution
        modelIrrev.c=zeros(nIrrevRxns,1);
        modelIrrev.c(objectiveColIrrev(i,1),1)=1;

        %find max objective
        FBAsolution = optimizeCbModel(modelIrrev);
        if (FBAsolution.stat ~= 1)
            not_solved=1;
            error('Failed to solve initial FBA problem');
        end
        maxObjective(i)=FBAsolution.f;
    end

    model2gimme = modelIrrev;
    model2gimme.c = zeros(nIrrevRxns,1);


    for i=1:nIrrevRxns
        if (expressionRxnsIrrev(i,1) > -1)   %if not absent reaction
            if (expressionRxnsIrrev(i,1) < threshold)
                model2gimme.c(i,1) = threshold-expressionRxnsIrrev(i,1); %FIX: use expression level as weight
            end
        end
    end

    for i=1:size(objectiveColIrrev,1)
        model2gimme.lb(objectiveColIrrev(i,1),1) = objectiveColIrrev(i,2) * maxObjective(i);
    end

    %gimmeSolution = optimizeCbModel(model2gimme,'min');
    
    % Replace optimizeCbModel with custom gurobi call and include integer cuts for ensemble solutions.
    
    nrgimme = length(model2gimme.lb);
    
    % Setting non-zero lower bounds
    lb = model2gimme.lb;
    ub = model2gimme.ub;
    for i = 1:nrgimme
        if lb(i) == 0
            lb(i) = 1e-6;
        end
        if ub(i) == 0
            ub(i) = -1e-6;
        end
    end
    
    %if N > 1
    %{
        Ycl = zeros(nrgimme,nRxns);
        Ycu = Ycl;
        for i = 1:nRxns
            for j = 1:length(rev2irrev{i})
                Ycl(rev2irrev{i}(j),i) = model2gimme.lb(rev2irrev{i}(j));
                Ycu(rev2irrev{i}(j),i) = model2gimme.ub(rev2irrev{i}(j));
            end
        end
    %}
    %end
    Ycl = zeros(nrgimme);
    Ycu = Ycl;
    Ycx = zeros(nRxns,nrgimme);
    for i = 1:nrgimme
        Ycl(i,i) = lb(i);
        Ycu(i,i) = ub(i);
    end
    for i = 1:nRxns
        if length(rev2irrev{i})>1
            Ycx(i,rev2irrev{i}(1)) = 1;
            Ycx(i,rev2irrev{i}(2)) = 1;
        end
    end
    Ycx = Ycx(any(Ycx,2),:);
    nyrev = size(Ycx,1);
    
    params.outputflag = 0;
    gmod = convertCobraModel2gurobi(model2gimme);
    gmod.modelsense = 'min';
    %if N > 1
        %gmod.A = [gmod.A,sparse(length(gmod.rhs),nRxns);speye(nrgimme),-sparse(Ycl);speye(nrgimme),-sparse(Ycu)];
        gmod.A = [gmod.A,sparse(length(gmod.rhs),nrgimme);speye(nrgimme),-sparse(Ycl);speye(nrgimme),-sparse(Ycu);sparse(nyrev,nrgimme),sparse(Ycx)];
        gmod.rhs = [gmod.rhs;zeros(2*nrgimme,1);ones(nyrev,1)];
        %gmod.lb = [gmod.lb;zeros(nRxns,1)];
        gmod.lb = [gmod.lb;zeros(nrgimme,1)];
        %gmod.ub = [gmod.ub;ones(nRxns,1)];
        gmod.ub = [gmod.ub;ones(nrgimme,1)];
        params.IntFeasTol = 1e-9;
        params.FeasibilityTol = 1e-9;
        params.OptimalityTol = 1e-6;
        s1 = '';
        s1(1:nrgimme) = '>';
        s2 = '';
        s2(1:nrgimme) = '<';
        s3 = '';
        s3(1:nyrev) = '<';
        vtype = '';
        %vtype(1:nRxns) = 'B';
        vtype(1:nrgimme) = 'I';
        gmod.sense = [gmod.sense,s1,s2,s3];
        gmod.vtype = [gmod.vtype,vtype];
        %gmod.obj = [gmod.obj;zeros(nRxns,1)];
        gmod.obj = [gmod.obj;zeros(nrgimme,1)];
        findex = length(gmod.sense);
        ysols = zeros(N,nrgimme);
        nActive = zeros(N,1);
    %end
    
    % Loop over number of expected solutions:
    keep_rxns = removed_reactions;
    for iter = 1:N
        t_init = cputime;
        msg = ['Generating model ',num2str(iter),' of ',num2str(N)];
        %disp(msg)
        waitbar(iter/N,f,msg);
        g1 = gmod;
        if iter > 1
            nsol = iter-1;
            g1.A = [g1.A;sparse(nsol,nrgimme),sparse(ysols(1:nsol,:))];
            %sm = sum(keep_rxns(:,1:nsol)',2);
            sm = nActive(1:nsol,:);
            sm = sm-1;
            g1.rhs = [g1.rhs;sm];
            g1.sense(findex+1:findex+nsol) = '<';
        end
        gimmeSol = gurobi(g1,params);
        gimme_objective(iter) = gimmeSol.objval;
        
        if ~ismember({gimmeSol.status},{'OPTIMAL'})%(gimmeSolution.stat ~= 1)
        %No solution for the problem
            display('Failed to solve GIMME problem');
            gimmeSol.x = zeros(nIrrevRxns,1);
            %gimmeSolution.x = zeros(nIrrevRxns,1);
        end
        
        reactionActivityIrrev = zeros(nIrrevRxns,1);
        curr_y = gimmeSol.x(nrgimme+1:end);
        for i=1:nIrrevRxns
            if curr_y(i) == 0
                reactionActivityIrrev(i,1) = 0;
            elseif ((expressionRxnsIrrev(i,1) > threshold) || (expressionRxnsIrrev(i,1) == -1))
                reactionActivityIrrev(i,1)=1;
            elseif (gimmeSol.x(i,1) > 0)
            %elseif (gimmeSolution.x(i,1) > 0)
                reactionActivityIrrev(i,1)=2;
            end
        end

        %Translate reactionActivity to reversible model
        reactionActivity = zeros(nRxns,1);
        for i=1:nRxns
            for j=1:size(rev2irrev{i,1},2)
                if (reactionActivityIrrev(rev2irrev{i,1}(1,j)) > reactionActivity(i,1))
                    reactionActivity(i,1) = reactionActivityIrrev(rev2irrev{i,1}(1,j));
                end
            end
        end
    
        remove = model.rxns(reactionActivity == 0);
        m1 = removeRxns(model,remove);
        interm_model = m1;
        %m1.lb(ismember(m1.rxns,remove)) = 0;
        %m1.ub(ismember(m1.rxns,remove)) = 0;
        
        [~,inactive] = FVA(m1);
        keepr = m1.rxns(~inactive);
        
        % Setting up integer cuts for next iteration
        
        
        for i = 1:nRxns
            if ~ismember(model.rxns(i),keepr)
                for j = 1:length(rev2irrev{i,1})
                    curr_y(rev2irrev{i,1}(1,j)) = 0;
                end
            end
        end
        
        ysols(iter,:) = curr_y';
        nActive = sum(ysols,2);
        
        removed_reactions(:,iter) = double(~ismember(model.rxns,keepr));
        keep_rxns(:,iter) = double(ismember(model.rxns,keepr));
        uT.Data(iter,1:5) = [iter gimme_objective(iter) sum(keep_rxns(:,iter)) (nRxns - sum(keep_rxns(:,iter))) (cputime - t_init)/60];
    end
    delete(f)
    delete(uT)
    delete(figx);
    
    
    %tissueModel = removeRxns(model,remove); 
end

function gmod = convertCobraModel2gurobi(model)

[nm,nr] = size(model.S);
gmod.A = sparse(model.S);
gmod.rhs = zeros(nm,1);
gmod.lb = model.lb;
gmod.ub = model.ub;
gmod.sense(1:nm) = '=';
gmod.vtype(1:nr) = 'C';
gmod.obj = model.c;
gmod.modelsense = 'max';

end
```
