## Supplementary material for "Guidelines for extracting biologically relevant context-specific metabolic models using gene expression data": iMAT_multi.m

```
function [removed_reactions,iMAT_objective] = iMAT_multi(model, expressionRxns, threshold_lb, threshold_ub, N, tol, core)
% Uses the iMAT algorithm (`Zur et al., 2010`) to extract a context
% specific model using data. iMAT algorithm find the optimal trade-off
% between inluding high-expression reactions and removing low-expression reactions.
%
% USAGE:
%
%    tissueModel = iMAT(model, expressionRxns, threshold_lb, threshold_ub)
%
% INPUTS:
%    model:             input model (COBRA model structure)
%    expressionRxns:    reaction expression, expression data corresponding to model.rxns.
%                       Note : If no gene-expression data are
%                       available for the reactions, set the
%                       value to -1
%    threshold_lb:      lower bound of expression threshold, reactions with
%                       expression below this value are "non-expressed"
%    threshold_ub:      upper bound of expression threshold, reactions with
%                       expression above this value are "expressed"
%
%
% OPTIONAL INPUTS:
%    tol:               minimum flux threshold for "expressed" reactions
%                       (default 1e-8)
%    core:              cell with reaction names (strings) that are manually put in
%                       the high confidence set (default - no core reactions)
%    logfile:           name of the file to save the MILP log (string)
%    runtime:           maximum solve time for the MILP (default value - 7200s)
%    epsilon:           value added/subtracted to upper/lower bounds
%                       (default 1)
%
% OUTPUT:
%    tissueModel:       extracted model
%
% `Zur et al. (2010). iMAT: an integrative metabolic analysis tool. Bioinformatics 26, 3140-3142.`
%
% .. Author: - Implementation adapted from the cobra toolbox
% (createTissueSpecificModel.m) by S. Opdam and A. Richelle, May 2017

f = waitbar(0,'Initializing iMAT runs');
figx = uifigure;
uT = uitable(figx);
uT.ColumnName = {'Model' 'Retained Reactions' 'Removed Reactions' 'Obj' 'Time (min)'}

if isfield(model,'C') || isfield(model,'E')
    issueConfirmationWarning('iMat does not handle the additional constraints and variables defined in the model structure (fields .C and .E.)\n It will only use the stoichiometry provided.');
end


%if nargin < 9 || isempty(epsilon)
    epsilon=1;
%end
%if nargin < 8 || isempty(runtime)
    %runtime = 7200;
    runtime = 7200;
%end
%if nargin < 7 || isempty(logfile)
%    logfile = 'MILPlog';
%end
if nargin < 6 || isempty(tol)
    tol = 1e-6;
end
if nargin < 7 || isempty(core)
    core={};
end
rxns = model.rxns;
[keeprxns,remove_rxns] = deal(zeros(size(model.S,2),N));
iMAT_objective = zeros(N,1);
nRxns = length(rxns);
    RHindex = find(expressionRxns >= threshold_ub);
    RLindex = find(expressionRxns >= 0 & expressionRxns < threshold_lb);
    
    %Manually add defined core reactions to the core
    if ~isempty(core)
        for i = 1:length(core)
            rloc = find(ismember(model.rxns, core{i}));
            if ~isempty(rloc) && isempty(intersect(RHindex,rloc))
                RHindex(end+1) = rloc;
            end
            if isempty(rloc)
                disp(['Manual added core reaction: ', core{i}, ' not found'])
            end
        end
    end

    S = model.S;
    lb = model.lb;
    ub = model.ub;
    nbasevar = size(S,2)+2*length(RHindex)+length(RLindex);
    nybase = 2*length(RHindex)+length(RLindex);

    % Creating A matrix
    A = sparse(size(S,1)+2*length(RHindex)+2*length(RLindex),2*size(S,2)+2*length(RHindex)+length(RLindex));
    [m,n,s] = find(S);
    for i = 1:length(m)
        A(m(i),n(i)) = s(i);
    end

    for i = 1:length(RHindex)
        A(i+size(S,1),RHindex(i)) = 1;
        A(i+size(S,1),i+size(S,2)) = lb(RHindex(i)) - epsilon;
        A(i+size(S,1)+length(RHindex),RHindex(i)) = 1;
        A(i+size(S,1)+length(RHindex),i+size(S,2)+length(RHindex)+length(RLindex)) = ub(RHindex(i)) + epsilon;
    end

    for i = 1:length(RLindex)
        A(i+size(S,1)+2*length(RHindex),RLindex(i)) = 1;
        A(i+size(S,1)+2*length(RHindex),i+size(S,2)+length(RHindex)) = lb(RLindex(i));
        A(i+size(S,1)+2*length(RHindex)+length(RLindex),RLindex(i)) = 1;
        A(i+size(S,1)+2*length(RHindex)+length(RLindex),i+size(S,2)+length(RHindex)) = ub(RLindex(i));
    end
    A1 = sparse([eye(size(S,2)),zeros(size(S,2),nybase),-diag(lb);eye(size(S,2)),zeros(size(S,2),nybase),-diag(ub)]);
    A = [A;A1];

    % Creating csense
    csense1(1:size(S,1)) = 'E';
    csense2(1:length(RHindex)) = 'G';
    csense3(1:length(RHindex)) = 'L';
    csense4(1:length(RLindex)) = 'G';
    csense5(1:length(RLindex)) = 'L';
    csense6(1:size(S,2)) = 'G';
    csense7(1:size(S,2)) = 'L';
    csense = [csense1 csense2 csense3 csense4 csense5 csense6 csense7];

    % Creating lb and ub
    lb_y = zeros(2*length(RHindex)+length(RLindex),1);
    ub_y = ones(2*length(RHindex)+length(RLindex),1);
    lb = [lb;lb_y;zeros(size(S,2),1)];
    ub = [ub;ub_y;ones(size(S,2),1)];

    % Creating c
    c_v = zeros(size(S,2),1);
    c_y = ones(2*length(RHindex)+length(RLindex),1);
    c = [c_v;c_y;zeros(size(S,2),1)];

    % Creating b
    b_s = zeros(size(S,1),1);
    lb_rh = lb(RHindex);
    ub_rh = ub(RHindex);
    lb_rl = lb(RLindex);
    ub_rl = ub(RLindex);
    b = [b_s;lb_rh;ub_rh;lb_rl;ub_rl;zeros(2*size(S,2),1)];

    % Creating vartype
    vartype1(1:size(S,2),1) = 'C';
    vartype2(1:2*length(RHindex)+length(RLindex),1) = 'B';
    vartype3(1:size(S,2),1) = 'B'; 
    vartype = [vartype1;vartype2;vartype3];

    for iter = 1:N
        t_init = cputime;
        waitbar(iter/N,f,['Extracting model ',num2str(iter),' of ',num2str(N)]);
        if iter > 1
            A1 = [keeprxns(:,1:iter-1)]';
            s1 = sum(A1,2);
            s1 = s1-1;
            Ax = [A;sparse(zeros(iter-1,nbasevar)),sparse(A1)];
            bx = [b;s1];
            csense8 = '';
            csense8(1:(iter-1)) = 'L';
            csensex = [csense csense8];
        else
            Ax = A;
            bx = b;
            csensex = csense;
        end
        MILPproblem.A = Ax;
        MILPproblem.b = bx;
        MILPproblem.c = c;
        MILPproblem.lb = lb;
        MILPproblem.ub = ub;
        MILPproblem.csense = csensex;
        MILPproblem.vartype = vartype;
        MILPproblem.osense = -1;
        MILPproblem.x0 = [];

        solution = solveCobraMILP(MILPproblem, 'timeLimit', runtime, 'printLevel', 3, 'relMipGapTol',0.005);

        x = solution.cont;
        rxnRemList = model.rxns(abs(x) < tol);
        keeprxns(:,iter) = double(~ismember(rxns,rxnRemList));
        iMAT_objective(iter) = solution.obj;
        uT.Data(iter,1:5) = [iter sum(keeprxns(:,iter)) (nRxns - sum(keeprxns(:,iter))) solution.obj (cputime - t_init)/60];
    end
    retained_reactions = keeprxns;
    removed_reactions = double(~logical(retained_reactions));
    delete(f)
    delete(uT)
    delete(figx);
    %tissueModel = removeRxns(model,rxnRemList); 
    
end
```
