## Supplementary material for "Guidelines for extracting biologically relevant context-specific metabolic models using gene expression data": checkModelFunction.m

```
function requiredMetsStatus = checkModelFunction(model, requiredMets)
% Checks that the model is able to produce the list of metabolites defined in
% `requiredMets`
%
% USAGE:
%    requiredMetsStatus = checkModelFunction(model, requiredMets)
%
% INPUTS:
%    model:                 model structure
%    requiredMets:          cell array with the list of metabolites that
%                           the model need to be able to produce
%
% OUTPUT:
%    requiredMetsStatus:    0 = test fail,
%                           1 = test succeed
%
% Authors: - This script is an adapted version of the implementation from
%            https://github.com/jaeddy/mcadre.
%          - Modified and commented by A. Richelle,May 2017
 

    if isempty(requiredMets) %If requiredMets is empty, skip the step and always say the test is passed
        requiredMetsStatus = true;
        %warning('No metabolites defined to check the model function')
    else
        % Add demand reactions for required metabolites
        % function evalc used to remove display of reaction added
        [~, model, requiredRxns] = evalc(['addDemandReaction(model,requiredMets);']);

        % Check that the demand reaction can carry fluxes (default value of
        % flux active - 1e-8)
        inactiveRequired = checkRxnFlux(model, requiredRxns);
        requiredMetsStatus = ~numel(inactiveRequired);
    end

end
```
