## Supplementary material for "Guidelines for extracting biologically relevant context-specific metabolic models using gene expression data": checkRxnFlux.m

```
function inactiveRequired = checkRxnFlux(model, requiredRxns)
% Checks that `requiredRxns` can carry fluxes
% This function uses the heuristic speed-up proposed by Jerby et al. in the MBA
% paper for performing a pseudo-FVA calculation.
%
% USAGE:
%   inactiveRequired = checkRxnFlux(model, requiredRxns)
%
% INPUTS:
%    model:                 model structure
%    requiredMets:          cell array with the list of rxns that
%                           the model need to have to be able to produce
%                           'requiredMets' (see checkModelFunction.m)
%
%OUTPUTS
%	inactiveRequired:       cell array with the list of rxns of
%                           'requiredRxns' that are inactives
%
% Authors: - This script is an adapted version of the implementation from
%            https://github.com/jaeddy/mcadre.
%          - Modified and commented A. Richelle,May 2017
 

    rxnList = requiredRxns;
    inactiveRequired = [];
    while numel(rxnList)
        
        numRxnList = numel(rxnList);
        model = changeObjective(model, rxnList);

        % Maximize all
        FBAsolution = optimizeCbModel(model, 'max');

        optMax = FBAsolution.x;

        % If no solution was achieved when trying to maximize all reactions, skip
        % the subsequent step of checking individual reactions
        if isempty(optMax)
            inactiveRequired = 1;
            break;
        end
        requiredFlux = optMax(ismember(model.rxns, requiredRxns));
        activeRequired = requiredRxns(abs(requiredFlux) >= 1e-8);
        rxnList = setdiff(rxnList, activeRequired);

        numRemoved = numRxnList - numel(rxnList);

        if ~numRemoved
            randInd = randperm(numel(rxnList));
            i = rxnList(randInd(1));
            model = changeObjective(model, i);

            % Maximize reaction i
            FBAsolution = optimizeCbModel(model, 'max');
            optMax = FBAsolution.f;
            if isempty(optMax)
                inactiveRequired = union(inactiveRequired, i);
                break;
            end
            if abs(optMax) < 1e-8
                inactiveRequired = union(inactiveRequired, i);
                break;
            end

            rxnList = setdiff(rxnList, i);
        end
    end
end
```
