## Supplementary material for "Guidelines for extracting biologically relevant context-specific metabolic models using gene expression data": connectivityEvidence.m

```
function E_C = connectivityEvidence(model, E_X)
% Computes the connectivity based evidence for non-core reactions using
% network topology
%
% USAGE:
%    E_C = connectivityEvidence(model, E_X)
%
% INPUTS:
%    model:    model structure
%    E_X:      expression-based evidence score
%
% OUTPUTS:
%    E_C:      connectivity-based evidence score
%
%
% Authors: - This script is an adapted version of the implementation from
%            https://github.com/jaeddy/mcadre.
%          - Modified and commented by S. Opdam and A. Richelle,May 2017

    Sbin = double(model.S ~= 0);
    % S matrix is binarized to indicate metabolite participation in each reaction

    % Adjacency matrix (i.e., binary reaction connectivity); the connection between
    % a reaction and itself is ignored by subtracting the identity matrix
    A = full(double(Sbin' * Sbin ~= 0));
    A = A - eye(size(A));

    % Influence matrix; describes the divided connectivity of reactions --
    % e.g., if R1 is connected to 4 reactions, its influence on each of those
    % reactions would be 0.25
    I = A ./ repmat(sum(A, 2), 1, size(A, 2));

    % Weighted influence matrix; each reaction's influence on others is
    % weighted by its expression score
    WI = repmat(E_X, 1, size(A, 2)) .* I;

    % Connectivity score; sum the influence of all connected reactions
    E_C = sum(WI)';
end
```
