## Supplementary material for "Guidelines for extracting biologically relevant context-specific metabolic models using gene expression data": pruningModel_v2.m

```
function [tissueModel, cRes] = pruningModel_v2(model, rankNonCore, coreRxn, zeroExpRxns, precursorMets, eta, tol,rmf,min_frac)
% Prunes the reactions of the model based on their expression,
% connectivity to core and confidence score.
%
% USAGE:
%   [tissueModel, cRes] = pruningModel(model, rankNonCore, coreRxn, zeroExpRxns, precursorMets, eta, tol)
%
% INPUTS:
%   model:               input model (COBRA model structure)
%   rankNonCore:         order for reaction removal
%	coreRxn:             core reactions in model
%	zeroExpRxns:         reactions with zero expression (i.e., measured zero, not just
%                        missing from expression data)
%   precursorMets:       list of metabolites involved in the
%                        reactions defined in protected reactions
%   eta:                 tradeoff between removing core and zero-expression
%                        reactions (default value: 1/3)
%   tol:                 minimum flux threshold for "expressed" reactions
%                        (default 1e-8)
%
% OUTPUTS:
%	tissueModel:         pruned, context-specific model
%	cRes:                result of model checks (consistency/function)
%                           - vs. +: reaction r removed from generic model or
%                           not
%                           1 vs. 2: reaction r had zero or non-zero expression evidence
%                           -x.y: removal of reaction r corresponded with removal of y (num.) total
%                           core reactions
%                           +x.1 vs. x.0: precursor production possible after removal of 
%                           reaction r or not
%                           3: removal of reaction r by itself prevented production of required
%                           metabolites (therefore was not removed)
%
%
% Authors: - This script is an adapted version of the implementation from
%            https://github.com/jaeddy/mcadre.
%          - Modified and commented by S. Opdam and A. Richelle,May 2017
 
    tissueModel = model;
    R_P = model.rxns;
    nonCoreRemoved = 0; 
    coreRemoved = 0;
    cRes = zeros(3000, 1);
    count = 1;
    
    % computing rmf_max
    rmf_max = zeros(size(rmf));
    for i = 1:numel(rmf_max)
        model = changeObjective(model,rmf(i),1);
        sol = optimizeCbModel(model);
        rmf_max(i) = sol.obj;
    end
    
    paramConsistency.epsilon=tol;
    paramConsistency.modeFlag=0;
    paramConsistency.method='fastcc';
    %fig1 = uifigure;
    %d1 = uiprogressdlg(fig1,'Title','Pruning Model using mCADRE','Message','Initializing...');
    n_noncore = numel(rankNonCore);
    while numel(rankNonCore) > 0
        
     %   display(['Reaction no. ', num2str(count)])
        r = rankNonCore(1);
        display(['Attempting to remove reaction ', r{:}, '...'])
        modelR = removeRxns(tissueModel, r);
        rmfst = false(size(rmf));
        for i = 1:numel(rmfst)
            modelR = changeObjective(modelR,rmf(i),1);
            sol = optimizeCbModel(modelR);
            if sol.obj >= rmf_max(i)*min_frac(i)
                rmfst(i) = true;
            else
                rmfst(i) = false;
            end
        end
        if sum(rmfst) == length(rmfst)
            rmf_stat = true;
        else
            rmf_stat = false;
        end

        % First check precursor production; if this test fails, no need to
        % check model consistency with FVA (time-saving step)
        rStatus = checkModelFunction(modelR, precursorMets);

        if rStatus && rmf_stat

            % Check for inactive reactions after removal of r
            if numel(r)
                % Remove reaction r from the model
                model_rem = removeRxns(tissueModel, r);
            end
            % Check for inactive reactions after removal of r
            [fluxConsistentMetBool,fluxConsistentRxnBool] = findFluxConsistentSubset(model_rem,paramConsistency,0);
            inactive_G= [ r; model_rem.rxns(fluxConsistentRxnBool==0)];
            
            inactiveCore = intersect(inactive_G, coreRxn);
            inactiveNonCore = setdiff(inactive_G, inactiveCore);

            % Remove reactions with zero expression (previously penalized in
            % rank_reactions) and corresponding inactive core reactions, only if
            % sufficiently more non-core reactions are removed
            if ismember(r, zeroExpRxns)
                
                display('Zero-expression evidence for reaction...')

                % Check model function with all inactive reactions removed
                modelTmp = removeRxns(tissueModel, inactive_G);
                tmpStatus = checkModelFunction(modelTmp, precursorMets);

                if (numel(inactiveCore) / numel(inactiveNonCore) <= eta) && tmpStatus
                    
                    R_P = setdiff(R_P, inactive_G);
                    tissueModel = removeRxns(tissueModel, inactive_G);

                    rankNonCore(ismember(rankNonCore, inactive_G)) = [];

                    nonCoreRemoved = nonCoreRemoved + numel(inactiveNonCore);
                    coreRemoved = coreRemoved + numel(inactiveCore);
                    num_removed = nonCoreRemoved + coreRemoved;
                    
                    display('Removed all inactive reactions')

                    % result = -1.x indicates that reaction r had zero
                    % expression evidence and was removed along with any
                    % consequently inactivated reactions; x indicates the number of
                    % core reactions removed
                    if numel(inactiveCore) > 100
                        removed_C_indicator = numel(inactiveCore) / 100;
                    else
                        removed_C_indicator = numel(inactiveCore) / 10;
                    end
                    result = -1 - removed_C_indicator;
                    
                else
                    % Note: no reactions (core or otherwise) are actually
                    % removed in this step, but it is necessary to update the
                    % total number of removed reactions to avoid errors below
                    num_removed = nonCoreRemoved + coreRemoved;
                    rankNonCore(1) = [];

                    display('No reactions removed')

                    % result = 1.x indicates that no reactions were removed
                    % because removal of r either led to a ratio of inactivated
                    % core vs. non-core reactions above the specified threshold
                    % eta (x = 1) or the removal of r and consequently
                    % inactivated reactions prevented production of required
                    % metabolites (x = 0)
                    result = 1 + tmpStatus / 10;
                end

            % If reaction has expression evidence, only attempt to remove
            % inactive non-core reactions
            else
                % Check model function with non-core inactive reactions removed
                modelTmp = removeRxns(tissueModel, inactiveNonCore);
                tmpStatus = checkModelFunction(modelTmp, precursorMets);

                if numel(inactiveCore) == 0 && tmpStatus
                    R_P = setdiff(R_P, inactiveNonCore);
                    tissueModel = removeRxns(tissueModel, inactiveNonCore);
         
                    rankNonCore(ismember(rankNonCore, inactiveNonCore)) = [];
 
                    nonCoreRemoved = nonCoreRemoved + numel(inactiveNonCore);
                    num_removed = nonCoreRemoved + coreRemoved;
                    display('Removed non-core inactive reactions')

                    % result = -2 indicates that reaction r had expression.
                    % evidence and was removed along with (only) non-core
                    % inactivated reactions; x indicates the number of
                    % core reactions removed (should be zero!)
                    if numel(inactiveCore) > 100
                        removed_C_indicator = numel(inactiveCore) / 100;
                    else
                        removed_C_indicator = numel(inactiveCore) / 10;
                    end
                    result = -2 - removed_C_indicator;
                    
                else
                    num_removed = nonCoreRemoved + coreRemoved;
                    rankNonCore(1) = [];

                    display('No reactions removed')

                    % result = 2.x indicates that no reactions were removed
                    % because removal of r either led to inactivated core
                    % reactions (x = 1) or the removal of r and consequently
                    % inactivated reactions prevented production of required
                    % metabolites (x = 0)
                    result = 2 + tmpStatus / 10;
                end
            end
        else
            num_removed = nonCoreRemoved + coreRemoved;
            rankNonCore(1) = [];

            % result = 3 indicates that no reactions were removed because
            % removal of r by itself prevented production of required
            % metabolites
            result = 3;
        end

        cRes(count) = result;
        count = count + 1;
        display(sprintf(['Num. removed: ', num2str(num_removed), ...
            ' (', num2str(coreRemoved), ' core, ', ...
            num2str(nonCoreRemoved), ' non-core); ', ...
            'Num. remaining: ', num2str(numel(rankNonCore)), '\n']))
        %d1.Value = 1 - numel(rankNonCore)/n_noncore;
        %d1.Message = [num2str(round(100*d1.Value,2)),'% Complete: ',num2str(num_removed),' reactions removed from the model']; 
    end
    %delete(d1);
    %delete(fig1);
    cRes(count:end) = [];
end
```
