## Supplementary material for "Guidelines for extracting biologically relevant context-specific metabolic models using gene expression data": rankReactions.m

```
function [coreRxn, nonCoreRxn, rankNonCore, zeroExpRxns] = rankReactions(model, ubiquityScore, confidenceScores, protectedRxns)
% Generate order for reaction removal.
% Gene ubiquity scores (reaction expression evidence) are used to
% define the core (`coreRxn`) and non-core (`nonCoreRxn`) reaction sets.
% Non-core reactions are ordered first by expression and then by
% connectivity evidence to give the list `P`. Any  reactions with zero
% expression are listed in `zeroExpRxns`.
%
% USAGE:
%    [coreRxn, nonCoreRxn, rankNonCore, zeroExpRxns] = rankReactions(model, ubiquityScore, confidenceScores, protectedRxns)
%
% INPUTS:
%    model:               input model (COBRA model structure)
%    ubiquityScore:       ubiquity scores corresponding to genes
%                         in `gene_id` quantify how often a gene is expressed accross samples.
%                         (default values defined in function of
%                         `threshold_high` value)
%    confidenceScores:    literature-based evidence for generic model,
%                         (default value = 0)
%    protectedRxns:       cell with reactions names that are manually added to
%                         the core reaction set (i.e. {'Biomass_reaction'})
%
% OUTPUTS:
%    tissueModel:         pruned, context-specific model
%    coreRxn:             core reactions in model
%    nonCoreRxn:          non-core reactions in model
%    rankNonCore:         order for reaction removal
%    zeroExpRxns:         reactions with zero expression (i.e., measured zero, not just
%                         missing from expression data)
%
%
% Authors: - This script is an adapted version of the implementation from
%            https://github.com/jaeddy/mcadre.
%          - Modified and commented by S. Opdam and A. Richelle,May 2017


    E_X=ubiquityScore;
    % Determine core reactions set from expression-based evidence
    % (includes reactions manually defined in
    % core + reactions having an expression > threshold_high)
    if ~isempty(protectedRxns)
        E_X(findRxnIDs(model,protectedRxns)) = 1;
    end
    coreRxn = model.rxns(E_X >=1);

    [nonCoreRxn, NC_idx] = setdiff(model.rxns, coreRxn);

    % Determine confidence level-based evidence
    E_L = confidenceScores;

    % Calculate connectivity-based evidence
    E_C = connectivityEvidence(model, E_X);

    % Rank non-core reactions
    E_X_NC = E_X(NC_idx); % expression-based evidence for non-core reactions
    E_C_NC = E_C(NC_idx); % connectivity-based evidence for non-core reactions
    E_L_NC = E_L(NC_idx); % literature-based evidence for non-core reactions
    [E_NC, NC_order] = sortrows([E_X_NC, E_C_NC, E_L_NC], [1, 2, 3]);
    NC_order = permute_NC(NC_order,E_NC);
    rankNonCore = nonCoreRxn(NC_order); % ordered (ranked) non-core reactions

    % Identify zero-expression reactions
    zeroExpRxns = rankNonCore(E_NC(:, 1) == -1e-6);
    
end

function NC_order = permute_NC(NC_order,E_NC)

x1 = unique(E_NC,'rows');
if isequal(size(x1,1),size(E_NC,1))
    disp('Model has no variability')
else
    for i = 1:size(x1,1)
        otemp = NC_order(ismember(E_NC,x1(i,:),'rows'));
        shf = randperm(length(otemp));
        shf = shf(:);
        NC_order(ismember(E_NC,x1(i,:),'rows')) = otemp(shf);
    end
end
    
    
    
end
    
```
