## Supplementary material for "Guidelines for extracting biologically relevant context-specific metabolic models using gene expression data": RMF_MBA_multi.m

```
function [removed_reactions] = RMF_MBA_multi(model, medium_set, high_set, N, tol,rmf_reaction)
% Uses the MBA algorithm (`Jerby et al., 2010`) to extract a context
% specific model using data. MBA algorithm defines high-confidence reactions
% to ensure activity in the extracted model. Medium confidence reactions
% are only kept when a certain parsimony trade-off is met. In random order,
% the algorithm prunes other reactions and removes them if not required to
% support high- or medium- confidence reactions.
%
% USAGE:
%
%    tissueModel = MBA(model, medium_set, high_set)
%
% INPUTS:
%    model:          input model (COBRA model structure)
%    medium_set:     list of reactions with medium confidence
%    high_set:       list of reactions with high confidence
%
% OPTIONAL INPUTS:
%    tol:            minimum flux threshold for "expressed" reactions
%                    (default 1e-8)
%
% OUTPUT:
%    tissueModel:    extracted model
%
% `Jerby et al. (201)). Computational reconstruction of tissue-specific
% metabolic models: application to human liver metabolism. Mol. Syst. Biol.
% 6, 401.`
%
% .. Author: - Commented by A. Richelle, May 2017.
% .. Adapted by Saratram Gopalakrishnan - March 2020 - Included the option to generate alternate solutions

if nargin < 5 || isempty(tol)
    tol = 1e-8;
end
f = waitbar(0,'Initializing MBA runs');
figx = uifigure;
uT = uitable(figx);
uT.ColumnName = {'Model' 'Retained Reactions' 'Removed Reactions' 'Time (min)'};
removed_reactions = zeros(length(model.rxns),N);
param.epsilon=tol;
    param.modeFlag=0;
    param.method='fastcc';
environment = getEnvironment();
nCore = 6;
oVar = 1:nCore:N;
model = changeObjective(model,rmf_reaction,1);
sol = optimizeCbModel(model);
gr_max = sol.obj;
for j = 1:length(oVar)
    imax = min(N+1-oVar(j),nCore);
    remR = zeros(size(removed_reactions,1),imax);
    tf = zeros(imax,1);
    msg1 = ['Extracting Model ',num2str(oVar(j)),' - ',num2str(oVar(j)+imax-1),' of ',num2str(N)];
    waitbar(oVar(j)/N,f,msg1);
    parfor mCount = 1:imax
        restoreEnvironment(environment);
        t_init = cputime;
        %msg1 = ['Extracting Model ',num2str(mCount),' of ',num2str(N)];
        %
        NC = setdiff(model.rxns,union(high_set,medium_set));
        nL = numel(NC);
        %MBA
        PM = model;
        removed = {};


        iter = 0;
        %disp(['extracting model ',num2str(mCount),' of ',num2str(N)])
        while ~isempty(NC)
            iter = iter+1;
            ri = randi([1,numel(NC)],1);
            r = NC{ri};       

            PM_temp = removeRxns(PM, r);
            sol = optimizeCbModel(PM_temp);
            if sol.stat ~= 1
                inactive = PM_temp.rxns;
            else
                [~,fluxConsistentRxnBool] = findFluxConsistentSubset(PM_temp,param,0);
                inactive=[PM_temp.rxns(fluxConsistentRxnBool==0);r];
            end  
            eH = intersect(inactive, high_set);
            eM = intersect(inactive, medium_set);
            eX = setdiff(inactive,union(high_set,medium_set));

            epsil=0.5;
            
            sol = optimizeCbModel(PM_temp);
            if isempty(sol.obj)
                sol.obj = 0;
            end
            if sol.obj >= 0.9*gr_max
                if numel(eH)==0 && numel(eM) < epsil*numel(eX)
                    PM = removeRxns(PM, inactive);
                    NC = setdiff(NC,inactive);
                    removed = union(removed,inactive);
                else
                    NC = setdiff(NC,r);
                end
            else
                NC = setdiff(NC,r);
            end
            %waitbar(1-numel(NC)/nL,f,[msg1,[': ',num2str(round(100*(1-numel(NC)/nL),2)),'%: ',num2str(numel(removed)),' Reactions Removed']]);
            disp([num2str(iter),' reactions evaluated. ',num2str(length(removed)),' of ',num2str(nL),' reactions removed']);
        end
        %waitbar(1-numel(NC)/nL,f,'Removing Reactions...');
        tissueModel = PM;
        %tissueModel = removeUnusedGenes(PM);
        tf(mCount) = (cputime - t_init)/60;

        remR(:,mCount) = double(~ismember(model.rxns,tissueModel.rxns));
    end
    removed_reactions(:,oVar(j):oVar(j)+imax-1) = remR;
    %tf = (cputime - t_init)/60;
    retR = length(remR(:,1))-sum(remR,1)';
    uT.Data(oVar(j):oVar(j)+imax-1,1:4) = [[oVar(j):oVar(j)+imax-1]' retR [sum(remR,1)]' tf];
    
end

delete(f)
delete(uT)
delete(figx);
    

end
```
