## Supplementary material for "Guidelines for extracting biologically relevant context-specific metabolic models using gene expression data": RMF_mCADRE.m

```
function [tissueModel] = RMF_mCADRE(model, ubiquityScore, confidenceScores, protectedRxns, checkFunctionality, eta, tol,rmf, rmf_frac)
% Uses the mCADRE algorithm (`Wang et al., 2012`) to extract a context
% specific model using expression data. mCADRE algorithm defines a set of core
% reactions and prunes all other reactions based on their expression,
% connectivity to core and confidence score. The removal of reactions is not
% necessary performed to support the core of defined functionalities. Core
% reactions are only removed if supported by a certain number of
% zero-expression reactions.
%
% USAGE:
%   tissueModel = mCADRE(model, ubiquityScore, confidenceScores)
%
% INPUTS:
%    model:                 input model (COBRA model structure)
%    ubiquityScore:         ubiquity scores corresponding to genes
%                           in gene_id quantify how often a gene is expressed accross samples.
%                           (default values defined in function of threshold_high value)
%    confidenceScores:      literature-based evidence for generic model
%                           (default value = 0)
%
% OPTIONAL INPUTS:
%    protectedRxns:         cell with reactions names that are manually added to
%                           the core reaction set (i.e. {'Biomass_reaction'})
%    checkFunctionality:    Boolean variable that determine if the model should be able
%                           to produce the metabolites associated with the protectedRxns
%
%                           0: don't use functionality check (default value)
%
%                           1: include functionality check
%    eta:                   tradeoff between removing core and zero-expression
%                           reactions (default value: 1/3)
%    tol:                   minimum flux threshold for "expressed" reactions
%                           (default 1e-8)
%
% OUTPUTS:
%    tissueModel:           pruned, context-specific model
%    coreRxn:               core reactions in model
%    nonCoreRxn:            non-core reactions in model
%    zeroExpRxns:           reactions with zero expression (i.e., measured zero, not just
%                           missing from expression data)
%    pruneTime:             total reaction pruning time
%    cRes:                  result of model checks (consistency/function)
%                           - vs. +: reaction `r` removed from generic model or not
%                           1 vs. 2: reaction `r` had zero or non-zero expression evidence
%                           `-x.y` : removal of reaction `r` corresponded with removal of `y` (num.) total
%                           core reactions
%                           `+x.1` vs. `x.0` : precursor production possible after removal of
%                           reaction `r` or not
%                           3: removal of reaction r by itself prevented production of required
%                           metabolites (therefore was not removed)
%
%
% `Wang et al. (2012). Reconstruction of genome-scale metabolic models for
% 126 human tissues using mCADRE. BMC Syst. Biol. 6, 153.`
%
% Authors: - This script is an adapted version of the implementation from
%            https://github.com/jaeddy/mcadre.
%          - Modified and commented by S. Opdam and A. Richelle,May 2017
 
    if isempty(protectedRxns)
        protectedRxns = {};
    end
    %if nargin < 5 || isempty(checkFunctionality)
    %    checkFunctionality=0; %don't use functionality check
    %end
    %if nargin < 6 || isempty(eta)
        eta = 1/3;% default value for tradeoff
    %end
    %if nargin < 7 || isempty(tol)
        %tol = 1e-8;% default tolerance
    %end
    
    model = changeObjective(model,rmf,1);
    sol = optimizeCbModel(model);
    rmf_max = sol.obj;

    %disp('Processing inputs and ranking reactions...')

    %[coreRxn, nonCoreRxn, rankNonCore, zeroExpRxns] = rankReactions(model, ubiquityScore, confidenceScores, protectedRxns);
    [coreRxn, ~, rankNonCore, zeroExpRxns] = rankReactions(model, ubiquityScore, confidenceScores, protectedRxns);

    % Precursor are defined as the metabolites involved in the
    % reactions defined in protected reactions
    %
    if checkFunctionality == 1 && ~isempty(protectedRxns)
        precursorMets = model.mets(any(model.S(:, findRxnIDs(model, protectedRxns)) < 0, 2));
        
        genericStatus = checkModelFunction(model, precursorMets);
        
    else
    	precursorMets={};
    	genericStatus=1;
    end
    %}
    if genericStatus
    	%disp('Generic model passed precursor metabolites test');

        %disp('Pruning reactions...')
   %     t0 = clock;
   %rem_rxns = prune_model(model,rankNonCore,coreRxn,zeroExpRxns,rmf_max,rmf_frac,eta);
   %tissueModel = removeRxns(model,rem_rxns);

        %[tissueModel, cRes] = pruningModel_v2(model, rankNonCore, coreRxn, zeroExpRxns, precursorMets, eta, tol, rmf, rmf_frac);
        tissueModel = pruningModel_v2(model, rankNonCore, coreRxn, zeroExpRxns, precursorMets, eta, tol, rmf, rmf_frac);
    %    pruneTime = etime(clock,t0);
    else
    	error('Generic model failed precursor metabolites test')
    end

    tissueModel = removeUnusedGenes(tissueModel);

end
```
