## Supplementary material for "Guidelines for extracting biologically relevant context-specific metabolic models using gene expression data": RMF_mCADRE_multi.m

```
function removed_reactions = RMF_mCADRE_multi(model,UbiquityScores,ConfidenceScores,N,prot_rxns,rmf_reaction)

f = waitbar(0,'Initializing mCADRE runs');
figx = uifigure;
uT = uitable(figx);
uT.ColumnName = {'Model' 'Retained Reactions' 'Removed Reactions' 'Time (min)'};

removed_reactions = zeros(length(model.rxns),N);
environment = getEnvironment();
nCore = 6;
oVar = 1:nCore:N;
done = false;
n_unique0 = 0;
j = 0;
ctr = 0;

model = changeObjective(model,rmf_reaction,1);
%sol = optimizeCbModel(model);
%gr_max = sol.obj;
if ~isempty(prot_rxns)
    prot_rxns = unique([prot_rxns;rmf_reaction]);
else
    prot_rxns = rmf_reaction;
end

while ~done
    j = j+1;
    imax = min(N+1-oVar(j),nCore);
    remR = zeros(size(removed_reactions,1),imax);
    tf = zeros(imax,1);
    msg1 = ['Extracting Model ',num2str(oVar(j)),' - ',num2str(oVar(j)+imax-1),' of ',num2str(N)];
    waitbar(oVar(j)/N,f,msg1);
    parfor i = 1:imax
        restoreEnvironment(environment);
        t_init = cputime;
        %waitbar(i/N,f,['Extracting model ',num2str(i),' of ',num2str(N)]);
        tmodel = RMF_mCADRE(model,UbiquityScores,ConfidenceScores,prot_rxns,0,1/3,1e-8,rmf_reaction,0.9);
        remR(:,i) = double(~ismember(model.rxns,tmodel.rxns));
        %uT.Data(i,1:4) = [i length(tmodel.rxns) sum(removed_Reactions(:,i)) (cputime - t_init)/60];
        tf(i) = (cputime - t_init)/60;

        remR(:,i) = double(~ismember(model.rxns,tmodel.rxns));
    end
    removed_reactions(:,oVar(j):oVar(j)+imax-1) = remR;
    %tf = (cputime - t_init)/60;
    retR = length(remR(:,1))-sum(remR,1)';
    uT.Data(oVar(j):oVar(j)+imax-1,1:4) = [[oVar(j):oVar(j)+imax-1]' retR [sum(remR,1)]' tf];
    U1 = unique(removed_reactions(:,1:(oVar(j)+imax-1))','rows');
    n_unique = size(U1,1);
    if n_unique > n_unique0
        n_unique0 = n_unique;
        ctr = 0;
        disp('New Solution Found')
    else
        ctr = ctr+1;
        disp('No new solution found')
    end
    if ctr >= 2000
        done = true;
    elseif j == length(oVar)
        done = true;
    end
end
removed_reactions = removed_reactions(:,1:(oVar(j)+imax-1));
delete(f)
delete(uT)
delete(figx);
end
    
```
