## Supplementary material for "Guidelines for extracting biologically relevant context-specific metabolic models using gene expression data": Supplementary Figure Captions.docx

**Supplementary Figure S1**

Box and Whisker plots showing the distribution of maximum flux relative to maximum flux predicted by the genome-scale model through the RMF reaction in models extracted for E. coli (A), 786O (B), and CHO-S (C) using GIMME, iMAT, MBA, and mCADRE with the global 60^th^ percentile (blue), global 75^th^ percentile (red), global 80^th^ percentile (yellow), StanDep (magenta), and Local T2 (green) thresholds.

**Supplementary Figure S2**

Receiver Operating Characteristics plots for models extracted using GIMME (blue), iMAT (orange), MBA (red), and mCADRE (magenta) for (A) *E. coli*, (B) 786O, and (c) CHO-S. The parent genome-scale model in each case is indicated using a green diamond marker.

**Supplementary Figure S3**

ROC plot for models extracted using GIMME (blue), iMAT ( orange), MBA (red), and mCADRE (cyan) for a lung cancer cell line (EKVX) using various thresholds. The parent genome-scale model is indicated using a black diamond marker.
