## Supplementary material for "Guidelines for extracting biologically relevant context-specific metabolic models using gene expression data": Supplementary Results.docx

**Model pre-processing prevents runtime errors**

Metabolic model pre-processing incorporates growth media information and measured uptake and secretion rates with the parent genome-scale metabolic model. Media composition is realized by limiting uptake by the model to only those metabolites present in the culture medium. This step inactivates reactions associated with carbon sources and trace minerals absent from growth media. When exometabolomic data (i.e., metabolite uptake and secretion rates) are available, corresponding lower and upper bounds of uptake and secretion reactions are specified in the model. The preprocessing step yields a flux-consistent model and sets of active or inactive reactions based on transcriptomics. For the models considered in this study, pre-processing removed 195, 307, and 355 reactions in *i*JO1366, Recon2.2, and *i*CHO1766, respectively, associated with highly expressed genes that are unable to carry flux. Because pruning methods can remove reactions only when all reactions associated with highly expressed genes are flux consistent, the inability of these reactions to carry flux causes the algorithm to falsely attribute flux inconsistency through these reactions to removal of non-core reactions. This leads to runtime errors which cause the algorithm to terminate without removing any reactions. Model pre-processing pre-emptively resolves such disagreements between the model and transcriptomics data that would otherwise lead to failures while extracting context-specific models.

**Model size and content are strongly influenced by the choice of extraction method**

Models extracted for *E. coli*, 786O, and CHO-S using GIMME, iMAT, MBA, and mCADRE showed substantial variation in model size and reaction content (Figure 3). In all three organisms, the smallest models were extracted using GIMME for all thresholds evaluated, whereas the largest models were extracted using MBA except for the Local T2 threshold. When the Local T2 threshold was applied, the largest models were generated using mCADRE.

Although the primary objective of the methods considered here is the maximal retention and flux consistency of reactions with highly expressed genes (i.e., core reactions) and/or the elimination of reactions with poorly expressed genes, differences in the combination and prioritization of objectives led to models of different sizes and contents. GIMME minimizes flux through all reactions below a specific threshold and therefore, attemptes to remove up to 1,694, 4,261, and 3,467 reactions in *E. coli*, 786O, and CHO-S, respectively, thus resulting in the smallest extracted models for all three organisms. On the other hand, MBA retained the greatest number of reactions (averaging 819, 1,871, and 1,341 reactions in *E. coli*, 786O, and CHO-S, respectively) resulting in the largest extracted models.

iMAT and mCADRE use fewer core reactions (averaging 572, 1,331, and 1,052 reactions in *E. coli*, 786O, and CHO-S, respectively) and generated models of intermediate sizes. Although both methods used the same set of core reactions, they differed in the stringency of retention of core reactions and the set of non-core reactions to be eliminated (Supplementary Table ST1). While mCADRE attempts to remove non-core reactions, iMAT only attempts to remove reactions with expression scores below the specified threshold (20^th^ percentile here). Unlike mCADRE, iMAT was more permissive to removing core reactions if at least an equal number of low-expression reactions were also removed. In contrast, mCADRE required at least twice as many non-core reactions to be removed to remove even one core reaction. Because of this, models generated using mCADRE were generally larger than those generated using iMAT.

Changing the threshold changed the number of protected core reactions for iMAT, MBA, and mCADRE, thereby contributing to variations in model sizes in each extracted ensemble. For global thresholding approaches, the number of core reactions decreased upon increasing the threshold for core reactions (Supplementary Table ST1, ST2, ST3). For example, the number of core reactions decreased from 536 to 225 reactions in *E. coli*, from 1,310 to 668 reactions in 786O, and from 940 to 381 reactions in CHO-S. The corresponding model sizes in ensembles generated using mCADRE decreased from 1,141 to 878 reactions in *E. coli* (Supplementary Figure S2A), from 2,510 to 1,749 reactions in 786O (Supplementary Figure S2B), and from 1,722 to 1,175 reactions in CHO-S (Supplementary Figure S2C).

While global thresholding approaches apply a uniform expression threshold for all genes in the model, the local T2 threshold applied individual gene-specific thresholds to determine whether reactions must be considered a core reaction. This caused the local T2 thresholding approach to be more inclusive of reactions, giving rise to a larger core reaction set (832 reactions in *E. coli*, 2,085 reactions in 786O, and 1,488 reactions in CHO-S) and larger models in all three organisms compared to the global thresholds.

Another approach, StanDep, performed a hierarchical clustering of gene expression data and applied cluster-specific thresholds. This allowed StanDep to identify housekeeping genes with constant and consistent expression and include those as core reactions. Thus, StanDep had the largest core reaction set (980 reactions in *E. coli*, 1,761 reactions in 786O, and 1,613 reactions in CHO-S) and the largest models of all evaluated thresholds (Figure 3). Interestingly, ensembles generated using GIMME showed minimal changes in model size in response to changes in the threshold. This is because GIMME only protects reactions coupled to the RMF reaction while maximally inactivating reactions with low expression scores. Since RMF-coupling is a structural property determined by the stoichiometric network, this set was unaffected by changes to expression thresholds, thereby resulting in model size-invariant ensembles.
