## Supplementary figures and images for "Guidelines for extracting biologically relevant context-specific metabolic models using gene expression data"

### Supplementary Figure S1.png

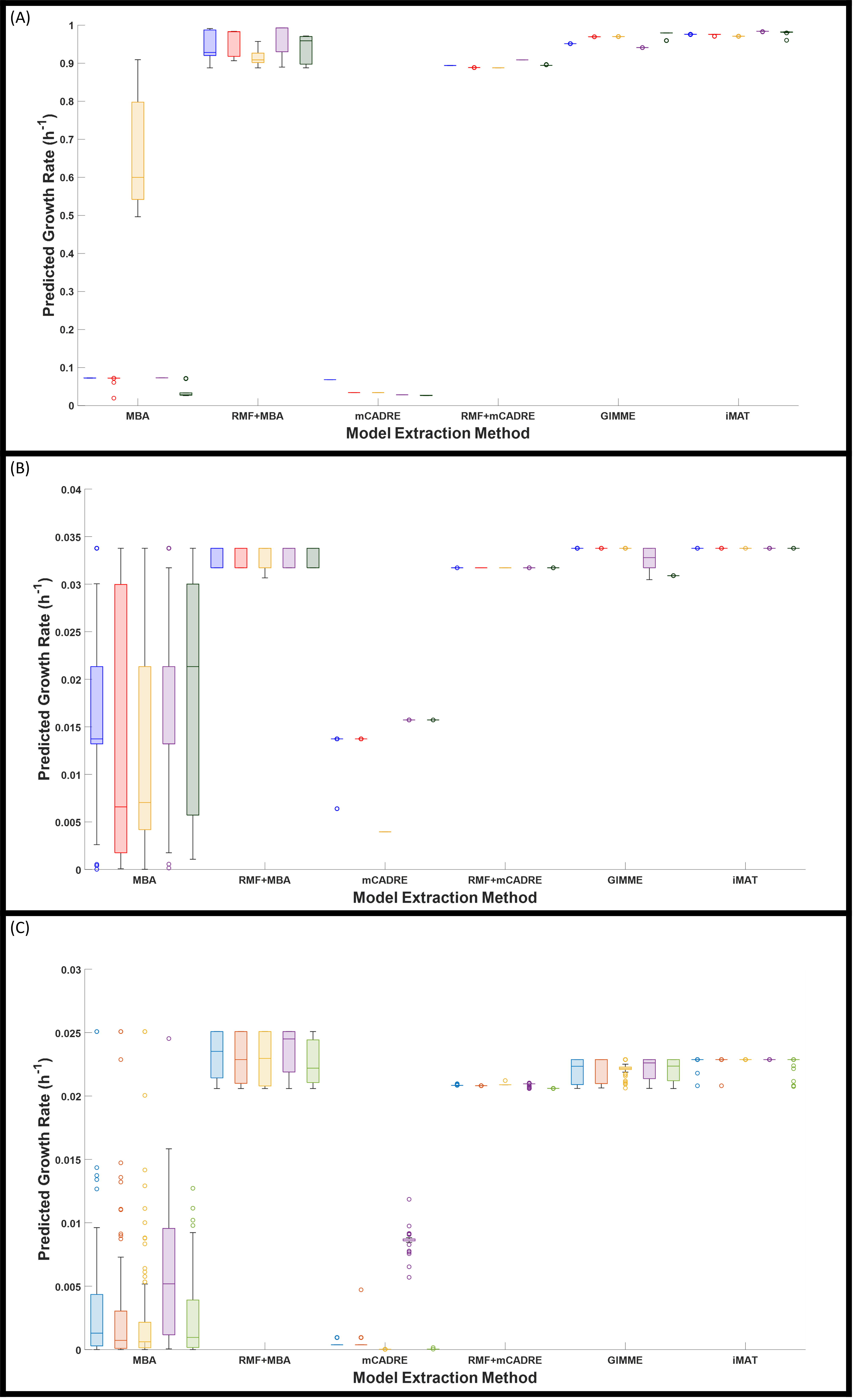

### Supplementary Figure S2.png

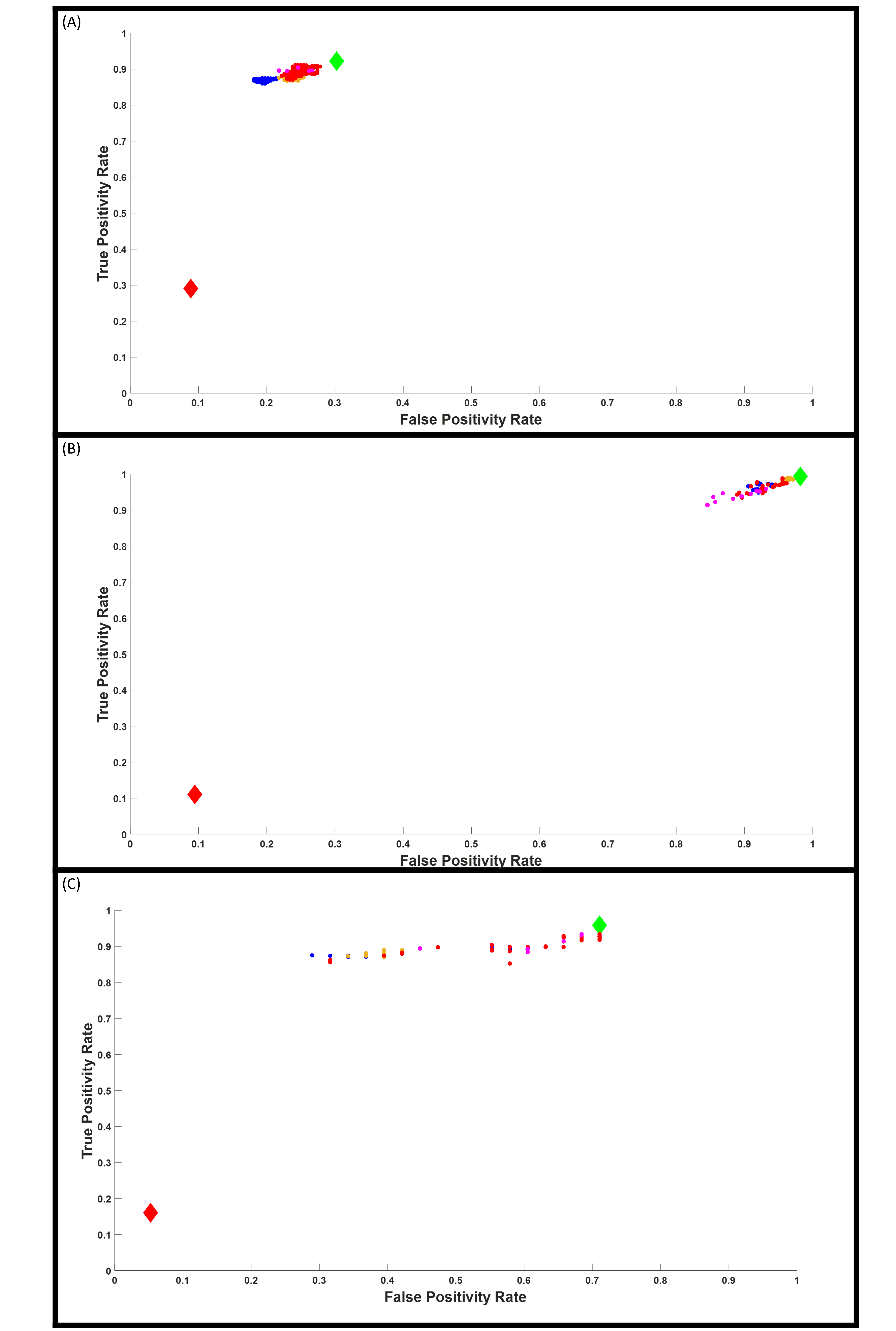

### Supplementary Figure S3.png

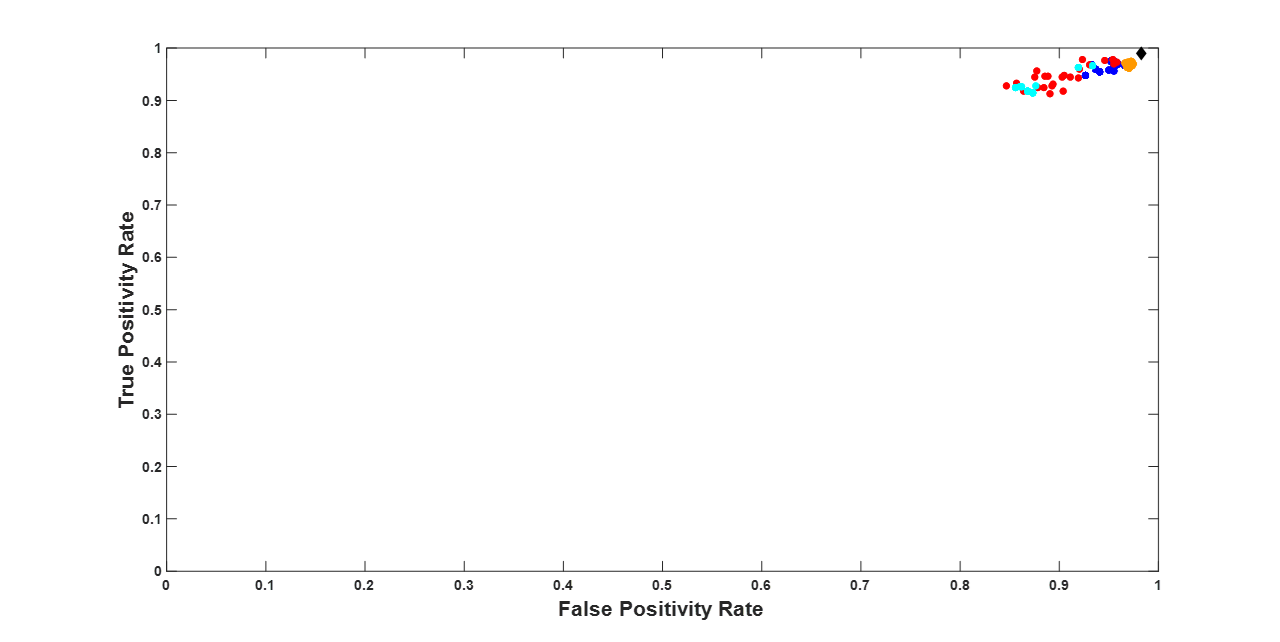
